# The interferon/STAT1 signaling axis is a common feature of tumor-initiating cells in breast cancer

**DOI:** 10.1101/2023.09.15.557958

**Authors:** Eric P. Souto, Ping Gong, John D. Landua, R. Rajaram Srinivasan, Abhinaya Ganesan, Lacey E. Dobrolecki, Stephen C. Purdy, Xingxin Pan, Michael Zeosky, Anna Chung, S. Stephen Yi, Heide L. Ford, Michael T. Lewis

**Affiliations:** Lester and Sue Smith Breast Center, Baylor College of Medicine, Houston, TX, 77030, USA; Cancer and Cell Biology Graduate Program, Baylor College of Medicine, Houston, TX, 77030, USA; Dan L Duncan Comprehensive Cancer Center, Baylor College of Medicine, Houston, TX, 77030, USA; Departments of Molecular and Cellular Biology and Radiology, Baylor College of Medicine, Houston, TX, 77030, USA; Department of Pharmacology, University of Colorado Anschutz Medical Campus (UC-AMC), Aurora, CO, 80045, USA; Pharmacology Graduate Program, UC-AMC, Aurora, CO, 80045, USA; University of Colorado Cancer Center, UC-AMC, Aurora, CO, 80045, USA; Livestrong Cancer Institutes and Department of Biomedical Engineering, The University of Texas at Austin, Austin, TX 78712, USA; Oden Institute for Computational Engineering and Sciences and Interdisciplinary Life Sciences Graduate Programs, The University of Texas at Austin, Austin, TX 78712, USA

## Abstract

A tumor cell subpopulation of tumor-initiating cells (TIC), or “cancer stem cells”, are associated with therapeutic resistance, as well as both local and distant recurrence. Enriched populations of TIC are identified by markers including aldehyde dehydrogenase (ALDH1) activity, the cell surface marker combination CD44^+^/CD24^−^, or fluorescent reporters for signaling pathways that regulate TIC function. We showed previously that <u>S</u>ignal <u>T</u>ransducer and <u>A</u>ctivator of <u>T</u>ranscription (STAT)-mediated transcription allows enrichment for TIC in claudin-low models of human triple-negative breast cancer using a STAT-responsive reporter. However, the molecular phenotypes of STAT TIC are not well understood, and there is no existing method to lineage-trace TIC as they undergo cell state changes. Using a <u>new</u> STAT-responsive lineage-tracing (LT) system in conjunction with our original reporter, we enriched for cells with enhanced mammosphere-forming potential in some, but not all, basal-like triple-negative breast cancer (TNBC) xenograft models (TNBC) indicating TIC-related and TIC-independent functions for STAT signaling. Single-cell RNA sequencing (scRNAseq) of reporter-tagged xenografts and clinical samples identified a common interferon (IFN)/STAT1-associated transcriptional state, previously linked to inflammation and macrophage differentiation, in TIC. Surprisingly, most of the genes we identified are not present in previously published TIC signatures derived using bulk RNA sequencing. Finally, we demonstrated that bone marrow stromal cell antigen 2 (BST2), is a cell surface marker of this state, and that it functionally regulates TIC frequency. These results suggest TIC may exploit the IFN/STAT1 signaling axis to promote their activity, and that targeting this pathway may help eliminate TIC.

**Significance:** TIC differentially express interferon response genes, which were not previously reported in bulk RNA sequencing-derived TIC signatures, highlighting the importance of coupling single-cell transcriptomics with enrichment to derive TIC signatures.

## INTRODUCTION

Breast cancer is a heterogeneous disease that can be classified into several histological and molecular subtypes (1,2). Triple-negative breast cancer (TNBC) is a histological subtype defined clinically by lack of estrogen receptor (ER) and/or progesterone receptor (PR), and without amplification or overexpression of the ERBB2 gene (HER2), and is an aggressive disease with poor outcomes (3).

There exists considerable intertumoral and intratumoral heterogeneity within each histological subtype. Intertumoral heterogeneity refers to variations across tumors, whereas intratumoral heterogeneity refers to the presence of molecularly distinct tumor cell populations within the same tumor. Both present major challenges clinically, particularly in TNBC, which itself represents at least six molecularly distinct subtypes (4,5). Due to the relative lack of targeted therapeutics, TNBC is typically treated with combination chemotherapies, with or without immuno-oncology agents or radiation.

With respect to intratumoral heterogeneity, there exists a subpopulation of cells, typically rare, termed tumor-initiating cells (TIC) (a.k.a. cancer stem cells (CSC)). TIC are defined functionally by their ability to self-renew, and to recapitulate clonally-derived cellular hierarchies present in the tumor from which they were isolated upon transplantation (6). Enriched TIC populations can be identified by aldehyde dehydrogenase (ALDH1) activity (7), the cell surface marker combination of CD44^+^/CD24^low/-^ (8), or fluorescent signaling reporters expressing pathway-specific transcription factors that regulate TIC function, including Wnt/β-catenin, Hedgehog, and Notch signaling, as well as POU5F1 (OCT4)/SOX2, and <u>S</u>ignal <u>T</u>ransducer and <u>A</u>ctivator of <u>T</u>ranscription 3 (STAT3)-mediated signaling (9–14). However, no single marker is present in all breast cancers.

Following neoadjuvant (before surgery) taxane– or anthracyclin-based chemotherapy in TNBC, residual breast cancer cells are enriched for TIC markers, including CD44^+^/CD24^low/-^ and aldehyde dehydrogenase (ALDH1) activity, and isolated cells show enhanced mammosphere-forming efficiency (MSFE), indicating that TIC are intrinsically resistant to some systemic therapies (15–18). Thus, standard cancer therapies can eliminate many “bulk” tumor cells to shrink tumors, but TIC survive preferentially, leading to local and/or distant recurrence. Indeed, some metastatic tumor cells express the TIC marker CD44, and such cells have enhanced tumor-initiating capacity (19). Thus, selective targeting of TIC, especially at metastatic sites, may be essential for effective treatment. Unfortunately, TIC-specific assays to evaluate drug efficacy are rarely done in clinical trials.

Here, we exploit a STAT signaling reporter, and a new STAT-responsive lineage-tracing (LT) system, to enrich cells with enhanced mammosphere-forming potential in some, but not all, non-claudin-low xenograft models of TNBC. Single-cell RNA sequencing (scRNAseq) on three xenograft models identified a distinct interferon (IFN)/STAT1-associated transcriptional state in TIC that is also present in most clinical samples evaluated, yet contains genes that have not been included in at least five TIC signatures derived through bulk RNA sequencing. Finally, we validated our signature by demonstrating that one of the genes, bone marrow stromal antigen 2 (BST2), can enrich for TIC, and regulates TIC frequency, but not self-renewal. Taken together, our data suggest that TIC exploit the IFN-mediated response pathway to regulate stem cell frequency independent of self-renewal. Thus, this pathway may represent an attractive candidate therapeutic target in a subset of breast cancers.

## MATERIALS & METHODS

### STAT lineage-tracing (LT) system construction

The STAT reporter (4M67-EGFP-P2A-CreER^T2^) was generated by LR Gateway® Cloning of a promoter donor vector and a gene donor vector into a destination vector. To generate a flexible promoter donor vector, the EF-1α promoter sequence from the L4/R1 EF-1α promoter donor vector (ThermoFisher Scientific, cat. A11146) was replaced with a 194bp sequence containing multiple restriction digestion sites to facilitate simple restriction cloning of any desired promoter. The STAT-specific promoter donor vector was generated by inserting the STAT signaling-specific 4M67 promoter from the STAT regulatory reporter (4M67-EGFP) (11) into the flexible promoter donor vector to generate the L4-4M67-R1 promoter donor vector. To generate the gene donor vector, a CreER^T2^ sequence with attB1/B2 recombination sites and an EcoRI cutting site between attB1 and CreER^T2^ was amplified from pCAG-CreER^T2^ (20) (a gift from Connie Cepko: Addgene plasmid #14797) by 2-step PCR, then incorporated into the pDonor233 vector (a kind gift from Dr. Kenneth L. Scott, Baylor College of Medicine, Houston, TX) with the P1/P2 compatible recombination sites by BP Gateway cloning. Next, GFP-P2A was amplified from pUltra (21) and inserted in front of CreER^T2^ using InFusion cloning to generate L1-EGFP-P2A-CreER^T2^-L2 gene donor vector. Gateway® Cloning was used to incorporate the 4M67 promoter donor vector and EGFP-P2A-CreER^T2^ gene donor vector into the pLenti6.4/R4R2/V5-Dest destination vector (ThermoFisher Scientific, cat. A11146), generating 4M67-EGFP-P2A-CreER^T2^.

To generate the dual color-switching lineage-tracing reporter (EFS-loxP-dsRed-loxP-Nep), the loxP-dsRed-loxP-GFP sequence with attB1/B2 recombination sites was amplified from pMSCV-loxP-dsRed-loxP-eGFP-Puro-WPRE (22) (a gift from Dr. Hans Clevers: Addgene plasmid #32702) by 2-step PCR and then incorporated into the pDonor233 vector with the P1/P2 compatible recombination sites by BP Gateway cloning. Gateway® Cloning was used to incorporate the EFS promoter donor vector and loxP-DsRed-loxP-EGFP gene donor vector into the pLenti6.4/R4R2/V5-Dest destination vector (ThermoFisher Scientific, cat. A11146). mNeptune2 was amplified from the mNeptune2-C1 vector (a kind gift from Dr. Michael Davidson; Addgene plasmid #54836) and used to replace the EGFP sequence by restriction cloning, generating EFS-_loxP_dsRed_loxP_-Nep.

### Lentivirus production and transduction

Lineage-tracing plasmids and envelope-expressing plasmids were prepared using a plasmid maxi kit (QIAGEN cat. 12162). To produce lentivirus, lineage-tracing plasmids and envelope-expressing plasmids were transfected into HEK293T cells using jetPRIME® transfection reagent (Polyplus, cat. 114-15). Lentiviral supernatants were collected every 24 hours for three days, then concentrated. Concentrated viruses were titered using the p24-Elisa kit (Takara Bio, cat. 632200) following manufacturer’s protocols. For lentiviral transduction, 5mg/mL polybrene (Santa Cruz Biotechnology, cat. sc-134220) was added to cell lines prior to lentiviral spinfection at 2500 RPM, 25°C for 30 minutes (MOI of 5). Transduced cells were incubated at 37°C, 5% CO_2_ for 24 hours, then lentivirus media was replaced with growth media. To generate LT cell lines (HepG2/C3A STAT LT, SUM159 STAT LT, and MDA-MB-231 STAT LT), 4M67-GFP-P2A-CreER^T2^ was transduced first, and transduced cells were selected using 10mg/mL blasticidin (Thermo Fisher Scientific, cat. A1113903). Following blasticidin selection, cell lines were then transduced with EFS-_loxP_dsRed_loxP_-Nep. Transduced cells were selected by FACS-enrichment of dsRed^+^ cells. To generate BST2 knockdown cell lines, SUM159 cells were lentiviral transduced with either one of two shRNA vectors (Horizon Discovery, sh-BST2 #1: cat. RHS3979-201824601, sh-BST2 #2: cat. RHS3979-201829090) or one non-targeting control vector (Horizon Discovery, cat. RHS6848) as described above. Transduced cells were selected using 1μg/mL puromycin (InvivoGen, cat. ant-pr-1).

Xenograft tumors were dissociated (isolation of single cells from human xenograft tumors). The resulting cells were transduced with the STAT regulatory reporter (4M67-GFP) (11) overnight in suspension in complete Mammary Epithelial Cell Growth Media (MEGM^+^) (Lonza, cat. CC-3153, with supplements, cat. CC-4136) containing 20ng/mL basic fibroblast growth factor (Invitrogen, cat. 13256-029), 10ng/mL epidermal growth factor (Invitrogen, cat. PHG0311), 4µg/mL Heparin (StemCell Technologies, cat. 07980), and 40 nL/mL B27 (Invitrogen, cat. 17504-044).

### Cell culture and *in vitro* treatment

HepG2/C3A, MDA-MB-231, and HEK293T cells were purchased from ATCC and were cultured under adherent conditions with phenol red-free DMEM (Gibco, cat. 31053-028), 10% charcoal-stripped FBS (Gibco, cat. 12-676-029), Penicillin-Streptomycin (GenDEPOT, cat. CA005-010), 1mM sodium pyruvate (Life Tech, cat. 11360070), and 2mM L-Glutamine (Thermo Fisher Scientific, cat. 25030081). SUM159 cells were purchased from BioIVT and were cultured under adherent conditions with phenol red-free Ham’s F12 (Caisson labs, cat. HFL05-500ML) containing 5% charcoal-stripped FBS, 5mg/mL insulin (Gemini Bioproducts, cat. 700-112P), 1mg/mL hydrocortisone (Sigma-Aldrich, cat. H-4001), and Penicillin-Streptomycin (GenDEPOT, cat. CA005-010). Cells were incubated in a 5% CO_2_ incubator at 37°C. All cell lines were authenticated using short tandem repeat profiling, which was performed by the Cytogenetics and Cell Line Authentication Core at MD Anderson Cancer Center (Houston, TX). Mycoplasma detection was routinely performed using Venor®GeM qOneStep Mycoplasma detection kit (Minerva Biolabs, cat. 11-91100). All experiments were performed with five passages of STR profiling and mycoplasma testing.

Human recombinant IL-6 (Sigma-Aldrich, cat. I1395) was prepared as a 50mg/mL stock solution in PBS/1%BSA. Recombinant human IFN alpha (IFNα) (Bio-Techne, cat. NBP2-35891-100ug) and recombinant human IFN gamma (IFNγ) (Bio-Techne, cat. 285-IF-100) were prepared as a 200mg/mL stock in PBS. Tamoxifen (TAM) (Sigma-Aldrich, cat. T5648) was prepared as a 5mM solution in 100% ethanol. HepG2/C3A STAT LT cells were treated with either IL-6 (50ng/mL), TAM (1µM), or the combination and the cells were visualized by confocal microscopy (Leica, TCS-SP5) and quantified by flow cytometry (BD LSRII analyzer) after 72 hours.

### Generation of cell line-derived and patient-derived xenografts

To prepare for cell injection, SUM159 STAT LT or MDA-MB-231 STAT LT cells were suspended in a 1:1 solution of their respective growth media and phenol red-free Matrigel® matrix (Corning, cat. 356237). Cells were transplanted into the #4 epithelium-free “cleared” fat pad of 3–4-week-old SCID/Beige mice (Charles River Laboratories, Wilmington, MA, http://www.criver.com). When palpable tumors formed, mice were administered TAM by intraperitoneal injection to induce dsRed to mNeptune2 color switching in STAT signaling (EGFP^+^) cells. TAM was prepared by dissolving in sunflower seed oil (Sigma-Aldrich, cat. S5007) and stirring overnight. All animal protocols were reviewed and approved by the Animal Protocol Review Committee at Baylor College of Medicine.

### Isolation of single cells from human xenograft tumors

Xenograft tumors were harvested and digested using 5mL digestion media (phenol red-free DMEM/F12 [Corning cat. 16-405-CV], 100mg/mL gentamicin [Worthington, cat. LS004176], 250U/mL collagenase II [Worthington cat. LS004174] and 2μg/mL DNase I [Stemcell, cat. 07900]) per 1g tumor at 37°C in a gentleMACS dissociator (Miltenyi). The tumor digestion was passed through a 200mM filter and centrifuged at 300g for 1 minute. Red blood cells were lysed by adding 9mL ddH_2_O to the cell pellet for 20 seconds and lysis was stopped by the addition of 1mL 10X PBS. To dissociate cell clumps further, the cell suspension was centrifuged at 300 g for 1 minute, then resuspended in TrypLE Express (Gibco, cat. 12605-010) for 5 minutes. To stop the digestion, 9mL of PBS was added, and the cell suspension was passed through a 40mm cell strainer.

### Analytical flow cytometry and FACS

Cells from adherent cultures were trypsinized and pelleted by centrifuging at 300g for 3 minutes. The cells were resuspended in HBSS^+^ (HBSS [Cytiva Life Sciences cat. SH30588.01], 2% FBS). Single cells from digested xenograft tumors were labeled using a BUV395 anti-mouse cocktail (pooled and diluted 1:40: anti-mouse H2kD [BD Biosciences, cat. 742437], anti-mouse CD45 [BD Biosciences, cat. 567451], anti-mouse CD31 [BD Biosciences, cat. 740231], and anti-mouse Ter119 [BD Biosciences, cat. 563827]) to exclude mouse cells. A far-red BST2 antibody was used to label BST2-expressing cells (diluted 1:40, BioLegend, cat. 348416). CD44 was labeled with a PE conjugated antibody (diluted 1:20, BD Biosciences, cat. 566803), and CD24 was labeled with a FITC conjugated antibody (diluted 1:20, BD Biosciences, cat. 655154). Cell viability was determined by Sytox Blue (diluted 1:1000, V450, Invitrogen, cat. S34857) staining. Cells were applied to a BD LSRII analyzer for analytical flow cytometry or a BD AriaII for fluorescence-activated cell sorting (FACS). Quantification of each cell population was performed using FlowJo (BD Biosciences) and statistical differences between cell populations were determined by one-way ANOVA followed by post-hoc Tukey’s test, which was calculated using GraphPad Prism version 9 (GraphPad Software, San Diego, CA, http://www.graphpad.com).

### ALDEFLUOR assay

ALDH activity was assessed using the ALDEFLUOR kit (StemCell technologies, cat. 01700) according to the manufacturer’s protocol. Briefly, cells were incubated in ALDEFLUOR assay buffer containing ALDH substrate for 30 minutes at 37°C. In each experiment, a sample of cells was stained under identical conditions with diethylaminobenzaldehyde (DEAB), a specific ALDH inhibitor, as a negative control. Any antibody incubation was performed immediately after ALDEFLUOR incubation, as described in the “analytical flow cytometry and FACS” section.

### Propidium Iodide cell cycle assay

Single cell suspensions were fixed in 70% ethanol overnight at –20°C. Then, cells were washed twice with PBS and stained in 100μL propidium iodide solution (0.001% Triton X-100, 0.2mg/mL DNase-free RNase A (Thermo Fisher, cat. 12091-021), 20μg/mL propidium iodide (Thermo Fisher, cat. P3566)) for 15 minutes at 37°C. After incubations, the cells were immediately applied to a BD LSRII analyzer for analytical flow cytometry.

### Immunohistochemistry and Immunofluorescence

Immunohistochemistry (IHC) and immunofluorescence (IF) were performed by the Lester and Sue Smith Breast Center Pathology Core. Tumors were fixed in 4% paraformaldehyde, paraffin-embedded, sectioned at 3–4 µm, and deparaffinized. The antigen retrieval buffer for pSTAT1, pSTAT3, and pSTAT5 staining was 0.1M Tris-HCL PH 9.0. Sections were then blocked in 3% hydrogen peroxide solution for 5 minutes and incubated by pSTAT1 Ser727 antibody (Cell Signaling Technology, cat. 8826, 1:200), pSTAT3 Y705 antibody (Cell Signaling Technology, cat. 9145, 1:25) or pSTAT5 antibody Tyr694 (Cell Signaling Technology, cat. 9359, 1:25), followed by Signalstain Boost IHC Detection Reagent (HRP, Rabbit) (Cell Signaling Technology, cat. 8114) for 30 minutes, visualized by 3, 3′ – Diaminobenzidine (DAB+) solution (DakoCytomation, Glostrup, Denmark), and counterstained by Harris hematoxylin. SUM159 tumors, known for activation of pSTAT3 (11), were used as a positive control for pSTAT3, and a human tissue array, which included the spleen, was used as a positive control for pSTAT1.

For IF, Five IF images were quantified for each PDX model using CellProfiler (23). 0.1M Tris-HCL PH 9.0. was used as the antigen retrieval buffer, then sections were blocked in 3% hydrogen peroxide solution for 5 minutes and incubated with pSTAT1 Ser727 antibody (Cell Signaling Technology, cat. 8826, 1:200), pSTAT3 Y705 antibody (Cell Signaling Technology, cat. 9145, 1:25), or a GFP antibody (Fisher Scientific, cat. NC9777966) overnight at 4°C. The following day, slides were incubated with secondary antibodies: anti-mouse (for pSTAT1 and pSTAT3, Thermo Fisher, cat. A-11032) and/or anti-rabbit (for GFP, Thermo Fisher, cat. A-11008) for one hour at room temperature. Slides were then counterstained with DAPI and imaged. For each PDX model, five representative images were quantified using FIJI (24).

### Mammosphere-forming efficiency assay

Following tumor digestion and FACS enrichment of fluorescent cell populations, equal numbers of cells from each fraction were plated in complete Mammary Epithelial Cell Growth Media (MEGM^+^) (Lonza, cat. CC-3153, with supplements, cat. CC-4136) containing 20ng/mL basic fibroblast growth factor (Invitrogen, cat. 13256-029), 10ng/mL epidermal growth factor (Invitrogen, cat. PHG0311), 4µg/mL Heparin (StemCell Technologies, cat. 07980), and 400nL/mL B27 (Invitrogen, cat. 17504-044) with 0.6% HPMC (Millipore Sigma, cat. M7027) to prevent cell aggregation, in an ultra-low attachment plate.

To prepare secondary mammospheres, primary mammospheres were collected and dissociated into single cells using TrypLE Express (Gibco, cat. 12605-010) for 5 minutes. To inactivate the TrypLE Express, 9mL of PBS was added, and the cell suspension was passed through a 40mm cell strainer. Then, an equal number of cells from each fraction were plated in MEGM^+^ media with 0.6% HPMC. Secondary mammospheres were imaged on a Yokogawa CV8000, and CellPathfinder (Yokogawa) was used to quantify the number of mammospheres in each well. Spheres with an area greater than 300mm^2^ and a diameter greater than 50mm were used to determine mammospheres. A BioTek Cytation 5 (Agilent, Santa Clara, CA) was used to take representative images. Statistical differences in mammosphere forming efficiency between cell populations were determined by one-way ANOVA followed by post-hoc Tukey’s test, which was calculated using GraphPad Prism version 9 (GraphPad Software, San Diego, CA, http://www.graphpad.com).

### Limiting-dilution transplantation assay

FACS-enriched cells were suspended in growth media at a concentration of 5,000 cells per 10μl (SUM159) or 50,000 cells per 10μl (BCM-4272) and serially diluted using growth media. Four dilutions were prepared (SUM159: 5,000, 1,000, 200, 40 cells per 10μl, BCM-4272: 50,000, 5,000, 500, 50 cells per 10μl), then phenol red-free Matrigel® matrix (Corning, cat. 356237) was added for a final 1:1 solution of growth media and Matrigel® before transplantation as described in “Generation of cell line-derived and patient-derived xenografts.” The mice were monitored until the first tumor reached 10% body weight (∼7-8 weeks) at which point the experiment was terminated and tumor outgrowth was recorded. Extreme limiting-dilution analysis was used to calculate the TIC frequency of each population (25).

### scRNAseq library prep and sequencing

SUM159, BCM-4272, and BCM-15006 were sorted based on fluorescent reporter activity (SUM159: EGFP^+^/mNeptune2^+^, EGFP^+^/dsRed^+^, EGFP^−^/mNeptune2^+^, EGFP^−^/dsRed^+^; BCM-4272 and BCM-15006: EGFP^+^ or EGFP^−^), and single-cell gene expression libraries were prepared according to Chromium Single Cell Gene Expression 3v3.1 kit (10x Genomics). In brief, single cells, reverse transcription (RT) reagents, Gel Beads containing barcoded oligonucleotides, and oil were loaded on a Chromium controller (10x Genomics) to generate single-cell GEMS (Gel Beads-In-Emulsions) where full-length cDNA was synthesized and barcoded for each single cell. Subsequently the GEMS are broken and cDNA from each single cell are pooled. Following cleanup using Dynabeads MyOne Silane Beads, cDNA is amplified by PCR. The amplified product is fragmented to optimal size before end-repair, A-tailing, and adaptor ligation. The final library was generated by amplification. Finally, single-cell sequencing libraries were sequenced using NovaSeq 6000 (read configuration: 28-10-10-90; 1% PhiX Spike-in; read depth 200M/sample).

For single nuclei sequencing, Tissue processing and nuclei RNA fixation from PDX samples were performed using Chromium Next GEM Single Cell Fixed RNA Sample Preparation Kit (10x Genomics). Fixation was then quenched and samples were stored at –80°C until sequencing. Single nuclei RNAseq libraries were prepared according to Chromium Single Cell Gene Expression 3v3.1 kit (10x Genomics), similar to the cDNA synthesis and barcoding process for each single cell. Single-cell nuclei RNA libraries were sequenced using NovaSeq (read configuration: Paired End 28-10-10-90; 1% PhiX Spike-in; read depth 100M/sample).

### scRNAseq data preprocessing and filtering

FASTQ files were mapped to the GRCh38 reference genome using the Cell Ranger Single Cell software v.6.1.2 (SUM159) or v7.0.1 (BCM-4272 and BCM-15006) (10X Genomics). Single nuclei sequencing data were integrated with the Texas Advanced Computing Center (TACC) supercomputer resources for data analysis and interpretation. In brief, gene expression counts for individual nuclei were generated from raw FASTQ files using Cell Ranger (version 3). Reads with low mapping quality (<10) or that map to multiple genomic locations were filtered out. Using Seurat v4.1.3 in R v4.2.2, the count matrix of each sample was used to create a Seurat object. Cells expressing greater than 1000 and less than 9000 genes. Cells with greater than 10% mitochondrial transcripts were excluded as they represented low-quality or dying cells. Next, fluorescent samples from the same tumor were merged into a single Seurat object (ex. BCM-4272 EGFP+ and BCM-4272 EGFP-merged to BCM-4272). To exclude the effects of cell cycle heterogeneity in the analysis, we used the CellCycleScoring function from Seurat. This function gives each cell a quantitative score for G1, G2/M, and S phases based on the expression of cell cycle phase genes (26). Subsequently, effects of cell cycle heterogeneity were regressed out of the data during data scaling.

### scRNAseq clustering and UMAP visualization

Seurat v4.1.3 was used for data normalization, dimensionality reduction, and clustering (27). The filtered gene matrices from each merged sample were normalized using the NormalizeData function. The normalized datasets were centered and scaled using ScaleData. Then, the most significant principal components were found using RunPCA, and ElbowPlot was used to determine the number of principal components to use for clustering each dataset. Different resolution parameters for unsupervised clustering were examined to find the optimal number of clusters based on clear segregation of clusters. For each dataset, the first 35 principal components were used for unsupervised clustering with a resolution = 0.1 for SUM159 (yielding nine clusters) and a resolution of 0.5 for BCM-4272 and BCM-15006 (yielding seven clusters and six clusters, respectively) using the FindNeighbors and FindClusters functions. To visualize the clustering for each dataset, the dimensionality was further reduced using the Seurat function RunUMAP. The same principal components used to determine the clustering were the same used to calculate the UMAP embedding. Each resulting cluster was further analyzed for potential doublets or low-quality cells using two methods: quality metrics (nCount_RNA, nFeature_RNA, and mitochondrial content) for each cluster were calculated and any outlier values (clusters with greater, or less than, two standard deviations than the average of all clusters) were removed. Three PDX models from an independent, single nuclei RNA seq, dataset were processed, clustered, and visualized as described above. UCell was used to score each cell based on similarity the 25 genes identified by intersecting SUM159-C7, 4272-C7, and 15006-C6 (28).

### scRNAseq differential gene expression analysis and intersection of gene lists

To examine the different transcriptional states of tumor cells, FindAllMarkers was used to identify differentially expressed genes in each cluster using the “bimod” likelihood-ratio test framework. Two additional parameters were used to include differentially expressed genes: genes that showed at least a 0.25 log2 fold expression difference, and genes that were expressed in at least 25% of the cells in a cluster. To compare the similarities between candidate TIC transcriptional states in different xenograft models, upregulated genes from each candidate TIC cluster were intersected using GeneOverlap (29). To examine the similarity between the TIC genes we identified and previously published TIC signatures derived using bulk RNA sequencing (30–34), we intersected the TIC genes common between any two xenograft models with previously published TIC signatures using GeneOverlap (29). To examine the extent to which the 25 common TIC genes are unique to a given transcriptional state in each xenograft model, we performed Marker Enrichment Modeling (MEM) using the 25 common TIC genes across models as MEM labels. Similarity scores based on MEM labels were calculated and visualized with a heatmap using the MEM_RMSD function (35).

### scRNAseq gene set enrichment analysis

To associate a biological phenotype with each cluster, gene set enrichment analysis was performed using ClusterProfiler (36). Differentially expressed genes from each cluster, as determine using the FindAllMarkers function described above, with a log fold change threshold of 0.25 were included in the analysis. Gene sets with a false discovery rate (FDR) < 0.01 and a Benjamini-Hochberg (BH)-adjusted p-value < 0.05 were considered significantly enriched, and these cutoffs were established prior to analyzing the data.

### Analysis of patient scRNAseq data

Publicly available human breast cancer datasets were retrieved through GEO (GSE161529, GSE176078) (37,38). Seurat v4.1.3 was used for cell cycle regression, data normalization, dimensionality reduction, and clustering, as described above. To identify clusters of epithelial cells, the expression of keratins (KRT8, KRT14, KRT18, and KRT19) was evaluated. Clusters with no keratin expression were filtered, and the epithelial dataset was re-normalized and clustered using Seurat v4.1.3. UCell was used to score every cell in each patient samples with more than 1,500 non-cycling epithelial cells based on similarity using the 25 genes identified by intersecting SUM159-C7, 4272-C7, and 15006-C6 (28). Harmony was used to integrate common cell states across patient samples (39). The patient samples were integrated by breast cancer subtype, pre-neoplastic, or normal breast. UCell was used to score each cell in the integrated datasets based on similarity using the 25 genes identified by intersecting SUM159-C7, 4272-C7, and 15006-C6 (28). A two-sided Wilcoxon test was used to determine significance of TIC scores. To identify differentially expressed genes between clusters, FindAllMarkers from Seurat v4.1.3 was used. To compare the similarities between clusters with high TIC scores across breast cancer subtypes and our 25-gene signature, upregulated genes from each cluster were intersected using GeneOverlap (29).

### Patient survival and chemotherapy response

To determine whether there is any association between our 25-gene TIC signature and overall survival in breast cancer patients, we used the RNAseq datasets from KM-plotter (40). ROC plotter was used to determine whether our 25-gene TIC signature is associated with chemotherapy response in breast cancer patients (41). Statistical significance was set at p < 0.05 before the analysis was performed.

### qRT-PCR

RNA extraction was performed using the RNeasy plus mini kit (Qiagen, cat. 74134) according to the manufacturer’s protocol. Next, we performed cDNA synthesis using SuperScript™ III first-strand synthesis system (ThermoFisher Scientific, cat. 18080051) according to the manufacturer’s protocol. Then, qPCR was performed using iTaq qPCR supermix (BioRad, cat. 1725130). Briefly, 10μL 2X Taq, 9μL cDNA, and 1μL of a 20X TaqMan probe (ThermoFisher Scientific, cat. Hs00171632_m1 (BST2), Hs00241666_m1 (ADAR), Hs00942643_m1 (OAS2), Hs00967084_m1 (PARP9), Hs00393816_m1 (PARP14), Hs01013996_m1 (STAT1), Hs99999901_s1 (18S)) were added per reaction. The qPCR reaction was performed in a C1000-Touch-Thermocycler (BioRad) (Initial denaturation: 95°C 30 seconds, followed by 39 cycles of denaturation: 95°C 5 seconds and annealing/extension 60°C 30 seconds). Samples were normalized to the expression of a housekeeping gene, 18S. Statistical differences in BST2 expression following chemotherapy were determined for each PDX independently by one-way ANOVA followed by post-hoc Tukey’s test. Statistical differences in the expression of IFN response genes were determined for each model independently by one-way ANOVA followed by post-hoc Tukey’s test. All statistical differences were calculated using GraphPad Prism version 9 (GraphPad Software, San Diego, CA, http://www.graphpad.com).

### Jess™ Automated Western Blot

BST2 expression levels in BST2 knockdown cells relative to the non-targeting control cells was evaluated using ProteinSimple’s Jess™ fully automated western blot analysis (Bio-Techne, cat. 004-650) according to the manufacturer’s protocol. A total of 0.5mg/mL protein was loaded per sample and a BST2 antibody (1:25, Novus Biologicals, cat. NBP2-27154) was used to detect BST2 levels in each sample. The ProteinSimple’s Jess™ protein normalization was used to normalize variation in protein loading.

## RESULTS

### STAT reporter activity is heterogenous in a cohort of triple-negative breast cancer PDX models

We showed previously that STAT-mediated transcriptional activity allows enrichment for TIC in two cell line xenograft (CDX) models of claudin-low breast cancer (SUM159 and MDA-MB-231) (11). While our focus was on STAT3 in these two models, STAT1 and STAT3 transcription factors bind similar DNA sequences, and both can activate the reporter (42).

To determine whether STAT signaling might also be associated with TIC function in a broader range of TNBC, we screened a cohort of 56 TNBC PDX models for pSTAT3 expression, of which 38 (68%) had detectable pSTAT3 staining in the epithelium (Supplementary Table 1). This value is higher than, but arguably consistent with, results using clinical samples of TNBC (∼37.5% (43)) given the known engraftment bias present in PDX collections toward immunologically “cold” tumors (44).

The 38 pSTAT3 positive TNBC models were then cross-referenced with PDX showing behaviors of interest with respect to putative roles of TIC, specifically local recurrence after apparent complete response to one or more chemotherapies, and/or metastasis to one or more organ sites (44,45). Based on these properties, a subset of 11 PDX models of TNBC were chosen for genetic tagging with the 4M67-EGFP lentiviral vector (11). Nine PDX showed varying degrees of EGFP expression, whereas candidate negative control models BCM-0046 and WHIM12, showed no detectable EGFP expression (Supplementary Table 2). Of nine EGFP+ PDX, five models (BCM-3611, BCM-4013, BCM-5097, HCI-028, and HCI-002) were excluded from further consideration because the percentage of EGFP^+^ cells was too low (<1.2%) to be experimentally tractable, thereby yielding four remaining PDX models for use.

The chosen PDX were then evaluated for pSTAT1, pSTAT3, and pSTAT5 by IHC, as well as EGFP expression by immunofluorescence (Fig. 1A, Fig. S1A). Consistent with pSTAT1/3 positivity by IHC, PDX models displayed their anticipated STAT reporter activity via detection of EGFP (Fig. 1A). Reporter-positive PDX models did not exhibit pSTAT5 staining, indicating STAT5 signaling is not responsible for reporter activity. In EGFP^+^ PDX, analytical flow cytometry (Fig. 1B) showed a continuum of EGFP reporter activity across models ranging from 7.88% (BCM-15006) to 31.7% (MC1) of tumor cells, ranges that should allow for discrimination of differential TIC function/gene expression in EGFP^+^ vs. EGFP^−^ cells, should such differences exist. Although EGFP percentage is comparable to pSTAT1 and pSTAT3 percentage (Fig. 1A), co-IF staining revealed little direct correlation between the reporter and pSTAT1/3 (Fig. S1B), likely due to the 26-hour half-life of GFP (46) relative to the comparatively rapid rate of STAT dephosphorylation (47).

**Figure 1:**
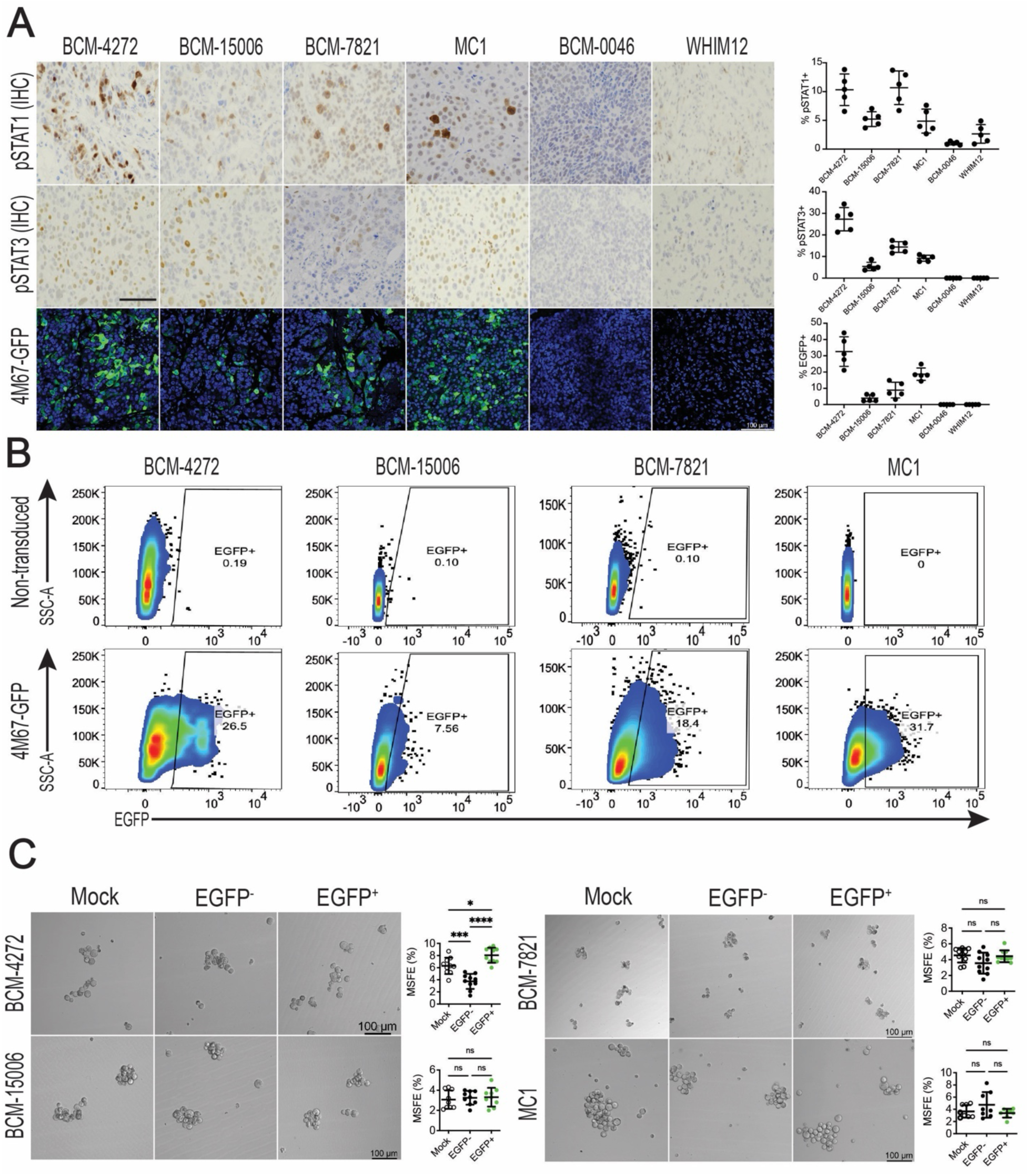
STAT reporter activity in breast PDX models. A) Representative IHC images showing levels of pSTAT1 and pSTAT3 in a panel of PDX models. 4M67-EGFP reporter activity was visualized by immunofluorescence (IF). Scale bar = 50 μm. IHC and IF were quantified using 5 representative images per PDX model. B) Flow cytometry plot illustrating the percentage of 4M67 reporter-positive cells in PDX models with demonstrated reporter activity. C) Representative images comparing the mammosphere forming efficiency between the EGFP^+^, EGFP^−^, and mock-sorted cells in four PDX models (BCM-4272: *n* = 10, BCM-15006: *n* = 9, BCM-7821: *n* = 10, MC1: *n* = 8) with a bar graph of the mammosphere forming efficiency. Values are mean ± SD. One-way ANOVA and Tukey’s post hoc test were used to determine statistical significance. *p < 0.05, ***p < 0.001, ****p < 0.0001. MSFE: mammosphere forming efficiency.

To determine whether STAT reporter activity enriches for cells with increased MSFE in reporter-positive TNBC PDX models (a surrogate for TIC function), we sorted EGFP^+^, EGFP^−^, and a mock sorted control from each reporter-positive model and plated them under low attachment conditions (Fig. 1C). STAT reporter positive cells in BCM-4272 had an 8.04% MSFE, representing a more than 100% increase relative to the EGFP^−^ cell population (3.74% MSFE), and the MSFE of the EGFP^−^ population was reduced by 59% relative to a mock sorted control (6.3% MSFE) (Fig. 1C). In contrast, STAT reporter activity did not enrich for cells with enhanced MSFE in BCM-15006, BCM-7821, or MC1 (Fig. 1C). Thus, while the 4M67 reporter can identify cells with enhanced MSFE in some TNBC, STAT-mediated transcription must have TIC-independent function(s).

### A STAT lineage-tracing reporter provides a long-term fluorescent label to STAT signaling cells

Given that TIC marker expression, including STAT transcriptional activity, is dynamically regulated (48), the inability to lineage-trace STAT signaling cells as they undergo cell state changes, or differentiate, is a major limitation of the 4M67-EGFP reporter.

To augment the 4M67-EGFP reporter, we developed a TAM-activated, Cre recombinase-dependent, STAT signaling-specific, lentiviral LT system (Fig. S2A–B). This system consists of two components: the first is a lentiviral vector that labels cells with active STAT signaling (EGFP^+^), followed by a self-cleaving P2A peptide and a TAM-activated Cre recombinase (4M67-EGFP-P2A-CreER^T2^) (Fig. S2A). The second component is a constitutively-expressed dual-color switching Cre-dependent reporter vector (EFS-_loxP_dsRed_loxP_-mNeptune2) (Fig. S2B). Addition of TAM drives color switching from red (dsRed) to far-red (mNeptune2) via CreER^T2^ recombination in STAT signaling cells within a specific temporal window where the TAM concentration remains high enough to activate Cre recombinase (6.8-hour half-life) (49) (Fig. 2A). Use of the LT system results in four cell populations: long-lived (label-retaining) STAT signaling cells (EGFP^+^/mNeptune2^+^), transient STAT signaling cells post-TAM (or cells that did not undergo Cre recombination) (EGFP^+^/dsRed^+^), lineage-tagged cells (EGFP^−^/mNeptune2^+^), and bulk tumor cells (EGFP^−^/dsRed^+^) (Fig. 2B).

**Figure 2:**
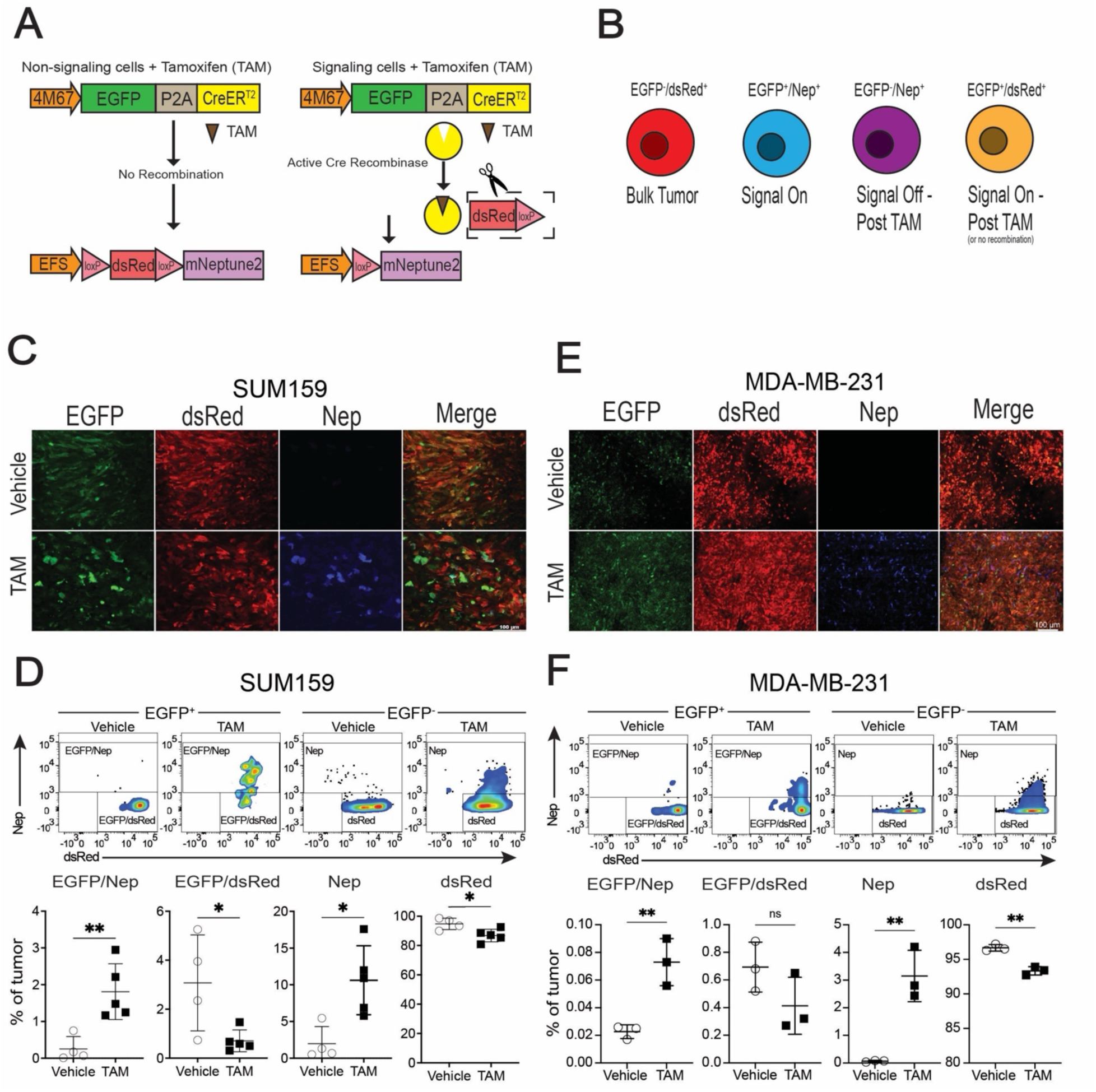
STAT lineage-tracing reporter provides a long-term fluorescent label to STAT signaling cells. A) Schematic illustrating how the STAT lineage-tracing system functions upon the addition of TAM. Cre is not transcribed in non-signaling (EGFP^−^) cells, thus no color-switching occurs in the presence of TAM. The Cre is transcribed in signaling (EGFP^+^) cells; therefore, the addition of TAM drives color-switching from dsRed to mNeptune2. B) Graphic illustrating the four different LT cell populations that can be identified by microscopy and flow cytometry. C) Confocal images of SUM159 STAT LT xenografts and D) flow cytometry quantification of the percent of each LT population two days after 4 mg TAM treatment (Untreated: *n* = 4, TAM-treated: *n* = 5). E) Confocal images of MDA-MB-231 STAT LT xenografts and F) flow cytometry quantification of the percent of each LT population two days after 4 mg TAM treatment (*n* = 3). One-way ANOVA and Tukey’s post hoc test were used to determine statistical significance. *p < 0.05, **p < 0.01, ns = not significant; values are mean ± SD. Nep: mNeptune2.

To demonstrate the sensitivity and specificity of the STAT LT reporter system, we used HepG2/C3A cells (IL-6 responsive) in a STAT3-dependent manner (11,50). Untreated STAT LT-transduced HepG2/C3A cells (HepG2/C3A STAT LT) had minimal EGFP and mNeptune2 expression. TAM (1µM) induced mNeptune2 expression in a small percentage of cells. Addition of IL-6 (50ng/mL) induced the EGFP reporter activity, without mNeptune2 activation, but the addition of both IL-6 and TAM induced EGFP and mNeptune2 expression (Fig. S2C–D). These results demonstrate the LT reporter system is both stringent and sensitive to STAT signaling, and TAM addition induced mNeptune2 expression in STAT-responsive cells.

To demonstrate the ability of the LT system to provide a long-term fluorescent label on STAT signaling cells *in vivo*, we introduced the LT reporters into the two TNBC cell lines used previously to demonstrate STAT reporter activity enriches for TIC (SUM159 STAT LT and MDA-MB-231 STAT LT) (11). To ensure effective lineage-tagging of STAT signaling cells, we performed an *in vivo* TAM dose-response analysis (Fig. 2SE–F). Given that high levels of TAM can cause toxicity in mice (51), we aimed to find a dose that resulted in maximal loxP recombination without any adverse effects. At 4mg, the recombination efficiency (the percentage of EGFP^+^/Nep^+^ cells out of all EGFP^+^ cells) two days after TAM treatment was 71%. The recombination efficiency, and the total percentage of mNeptune2^+^ cells, decreased with each successive dose, with the lowest dose (0.006mg) being comparable to vehicle. Given high recombination with no observable toxicity at 4mg TAM, we used this dose for the remainder of our studies.

SUM159 STAT LT or MDA-MB-231 STAT LT cell lines were engrafted into SCID/beige mice, and tumor-bearing mice were treated with 4mg TAM when tumors reached 100mm^3^. Two days after TAM treatment, we visualized all four LT cell populations in situ using confocal microscopy and quantified their abundance in each tumor model using flow cytometry (Fig. 2C–F). In both cell line xenografts, TAM treatment increased the percentage of EGFP^+^/Nep^+^ cells (from 0.25% to 1.8% in SUM159, and from 0.02% to 0.07% in MDA-MB-231), which was accompanied by a decrease in EGFP^+^/dsRed^+^ cells (from 3.08% to 0.7% in SUM159 from 0.7% to 0.4% in MDA-MB-231). Likewise, TAM treatment increased the percentage of mNeptune2^+^ cells (from 1.98% to 8.88% in SUM159 and from 0.07% to 3.15% in MDA-MB-231) with a corresponding decrease in the percentage of dsRed^+^ cells (from 94.7% to 88.3% in SUM159 and from 96.7% to 93.3% in MDA-MB-231). These results indicate that TAM treatment leads to Cre-dependent mNeptune2 expression *in vivo*, demonstrating the utility of the STAT LT system to study STAT TIC biology.

### Long-lived STAT signaling cells in SUM159 xenografts have enhanced mammosphere-forming efficiency

To confirm that the STAT LT system enriches for cells with enhanced MSFE from SUM159 tumors in a manner similar to that of the 4M67-EGFP vector shown previously (11), we dissociated SUM159 xenografts eight days after TAM treatment, sorted the four fluorescent cell populations (EGFP^+^/mNeptune2^+^, EGFP^+^/dsRed^+^, EGFP^−^/mNeptune2^+^, and EGFP^−^/dsRed^+^), and performed MSFE assays (Fig. 3A–B). Long-lived, label-retaining STAT signaling cells (EGFP^+^/mNeptune2^+^) had a significantly higher MSFE (7.2%) compared to all other cell populations, which were not statistically different from each other.

**Figure 3:**
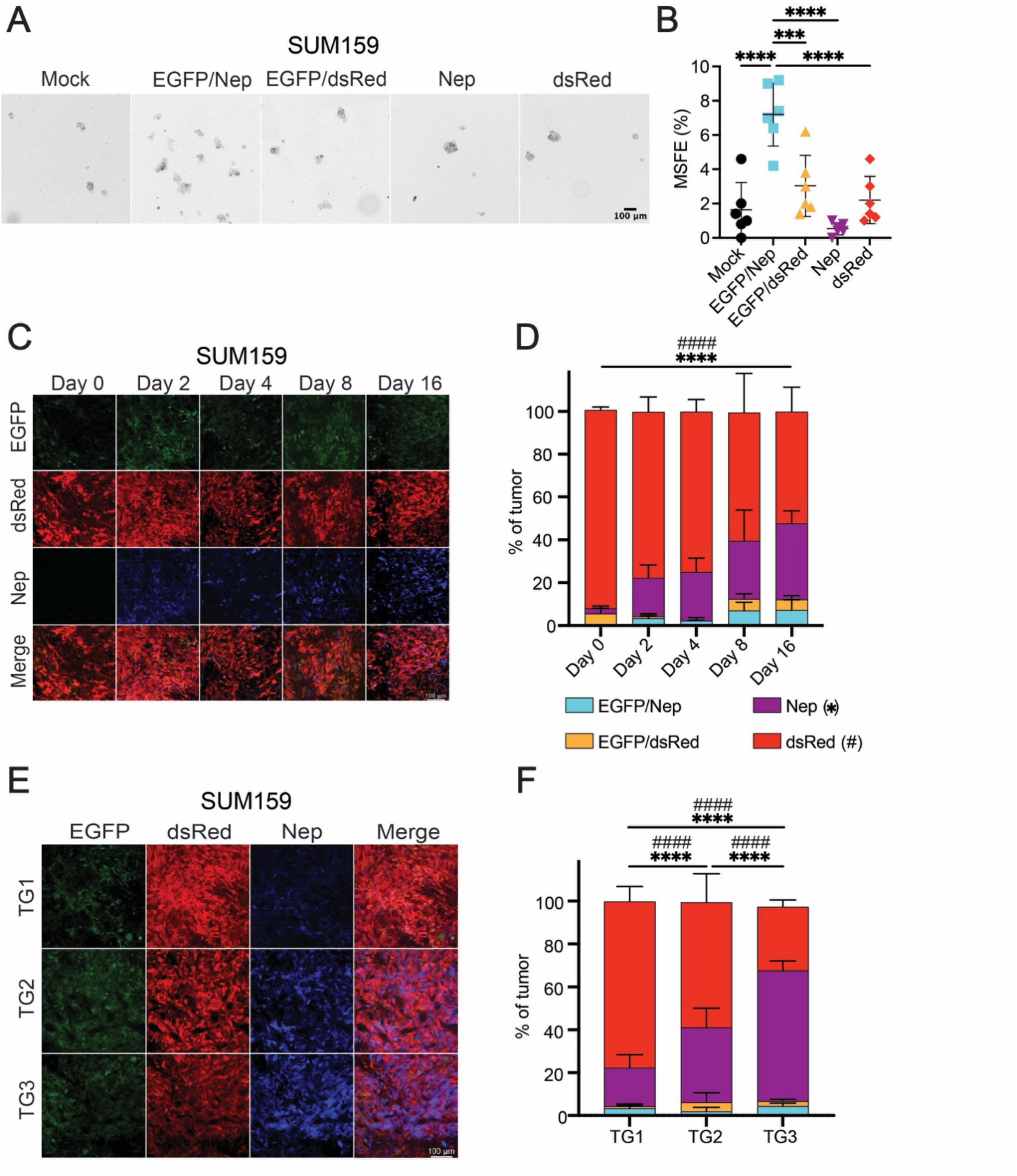
Long-lived STAT signaling cells are enriched for TIC in SUM159 xenografts. A) SUM159 STAT LT xenografts were dissociated, then each LT population was FACS enriched and grown as mammospheres. B) Mammosphere forming efficiency of each LT cell population from SUM159 xenografts (*n* = 6). One-way ANOVA and Tukey’s post hoc test were used to determine statistical significance. C) Confocal images of a 16-day TAM time course in SUM159 xenografts (*n* = 6). D) Flow cytometry quantification of the percentage of each LT cell population in SUM159 xenografts at five different time points. E) The time course was extended using a serial transplantation assay. Tumor fragments harvested two days post TAM (TG1), were transplanted into new hosts two additional times (*n* = 6). F) Flow cytometry quantification of the percentage of each LT cell population in SUM159 xenografts at each transplant generation (TG). ***p < 0.001, ****p < 0.0001, ^####^p < 0.0001; values are mean ± SD. Nep: mNeptune2.

The cancer stem cell hypothesis postulates that cellular heterogeneity in tumors results, in part, from hierarchical organization of cell types present, and that these hierarchies are fueled by rare, self-renewing TIC (52). We reasoned that long-lived STAT signaling cells give rise to progenitor cells (EGFP^−^/mNeptune2^+^) that will differentiate into non-TIC bulk tumor cells over time.

To test this hypothesis, we performed a 16-day time course study. We harvested SUM159 tumors prior to TAM treatment, and again at days 2, 4, 8, and 16 after the TAM pulse (Fig. 3C–D). A total of 74% of the EGFP^+^ cells recombined (EGFP^+^/dsRed^+^ converted to EGFP^+^/mNeptune2^+^ cells) two days after the TAM pulse. Between days 0 and 16, the percentage of EGFP^−^/Nep^+^ cells increased from 2% to 35% of the tumor (p < 0.001), indicating lineage-tagged cells proliferate and displace the EGFP^−^/dsRed^+^ cells. Furthermore, we noted the re-emergence of EGFP^+^/dsRed^+^ cells on the 8^th^ day, suggesting STAT signaling activation in cells not defined by STAT signaling at the time of the TAM pulse.

To extend the timeframe of this experiment, we serially transplanted tumor fragments two days after the TAM pulse for a total of three transplant generations (TG) (Fig. 3E–F). Each TG was allowed to grow for 21 days and harvested. The percentage of lineage-tagged cells increased from 18% to 35% from TG1 to TG2, then to 61% in TG3. Furthermore, the total percentage of EGFP^+^/mNeptune2^+^ cells remained unchanged over time and following serial passaging, indicating tight regulation of STAT signaling and TIC frequency.

### Interferon response genes are differentially expressed in TIC

To determine whether STAT reporter-positive cells express a distinct set of genes that may regulate TIC biology, we FACS-enriched all reporter populations from three TNBC xenograft models: two models where STAT reporter activity enriched for TIC (SUM159 and BCM-4272), and one model where STAT reporter activity did not enrich for TIC (BCM-15006), and performed scRNAseq (Fig. 4A).

**Figure 4:**
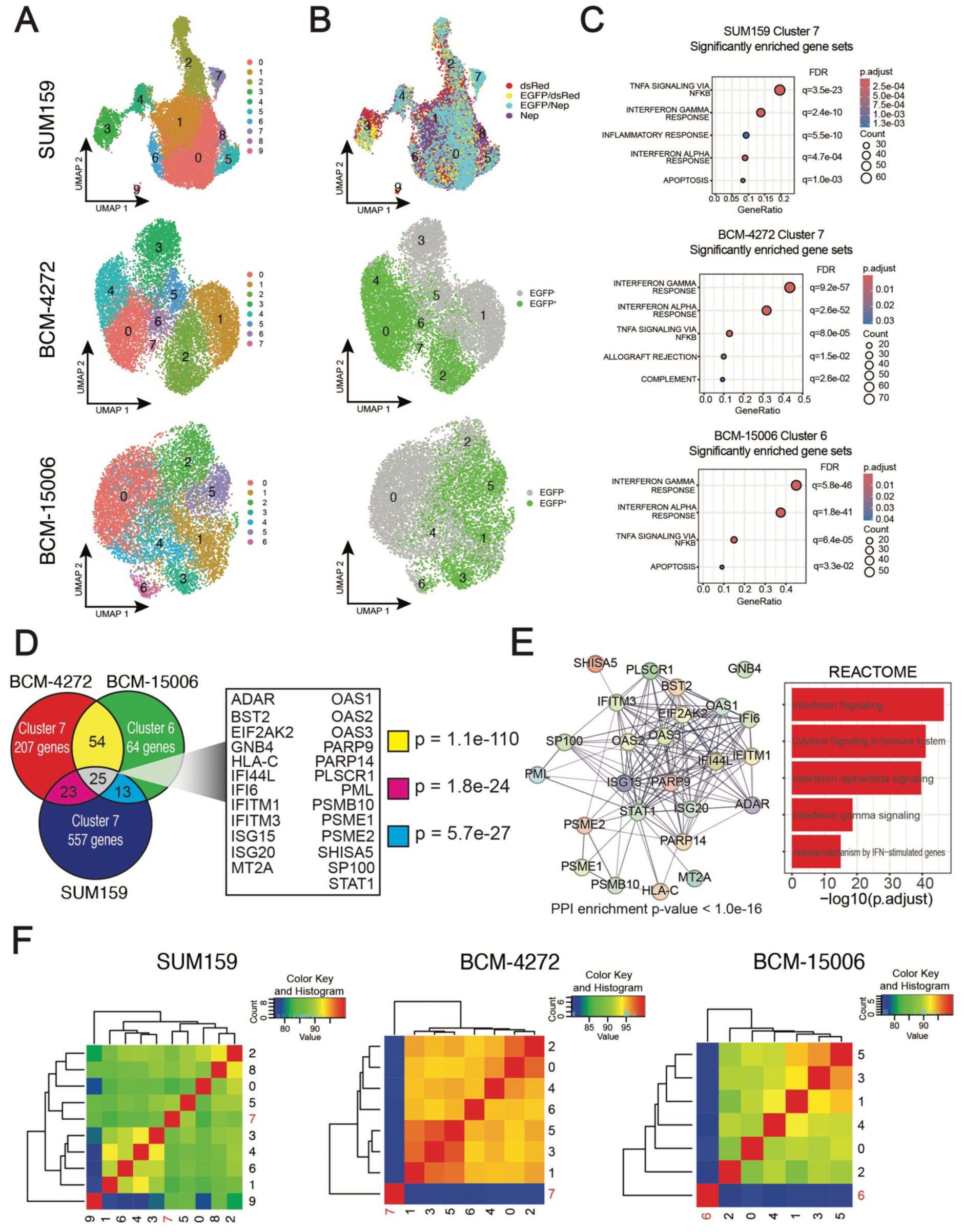
Interferon response genes are differentially expressed in TIC. SUM159 STAT3 LT, BCM-4272 4M67-EGFP, and BCM-15006 4M67-EGFP xenografts were harvested and dissociated, then each fluorescent population was enriched and sequenced using 10X genomics. A) Each xenograft was clustered and visualized using a UMAP plot. B) UMAP with cell identity overlayed on the plot. C) Hallmark gene set enrichment analysis of candidate TIC transcriptional state from each xenograft. D) Intersection of the upregulated genes in candidate TIC clusters with accompanying pairwise comparisons using Fisher’s exact test. Intersection of candidate TIC clusters from each model reveals 25 shared genes. E) STRING database illustration of protein-protein interactions between the 25-gene TIC signature and overrepresentation analysis results using the 25-gene list. F) Marker enrichment modeling heatmap illustrating cluster-to-cluster similarity based on expression of the genes in the 25-gene TIC signature. Nep: mNeptune2, FDR: false discovery rate, p.adjust: Benjamini-Hochberg (BH)-adjusted p-values.

To determine whether EGFP reporter activity could be used to identify TIC transcriptional states in each model, we overlayed sample information on Uniform Manifold Approximation and Projection (UMAP) plots to visualize whether reporter positive cells clustered together (Fig. 4B, Fig. S3B–D). Then, we looked at the percent representation of each cluster within each reporter population to identify clusters that were overrepresented in EGFP positive samples. In SUM159, though all clusters are represented in the EGFP^+^ cells, positive cells are highly overrepresented in cluster 7 (SUM159-C7) (9.8% of the EGFP^+^/mNeptune2^+^, 8.2% of the EGFP^+^/dsRed^+^ cells, vs. 0.1% of mNeptune2^+^ cells, and 0.07% of dsRed^+^ cells) (Fig. 4B, Fig S3B), suggesting it represents a TIC transcriptional state. In BCM-4272, clusters 0 (35% of EGFP^+^ cells, 0.5% of EGFP^−^ cells), 2 (25% of EGFP^+^ cells, 10% of EGFP^−^ cells), 4 (22% of EGFP^+^ cells, 0.7% of EGFP^−^ cells), and 7 (0.5% of EGFP^+^ cells, 0.1% of EGFP^−^ cells) are highly represented in the EGFP^+^ cells, suggesting at least one of these clusters may represent a TIC transcriptional state (Fig. 4B, Fig. S3A).

To clarify which of these clusters most likely represents a TIC transcriptional state, we performed differential gene expression analysis between these four clusters and found that cluster 7 differentially expressed genes associated with TIC, including *ABCD1*, *CXCL11*, *NOTCH1*, *RELA*, and *TCF7*, suggesting cluster 7 (4272-C7) represents TIC (Fig. S3B).

Because STAT reporter activity did not enrich TIC (Fig. 1C–D) in BCM-15006, we reasoned that a cluster representing a TIC transcriptional state would not be overrepresented in either EGFP^+^ and EGFP^−^ cells. Cluster 6 (15006-C6) is the only cluster that was equally represented in both EGFP^+^ and EGFP^−^ cells (2.1% of EGFP^+^, 1.9% of EGFP^−^ cells), suggesting this cluster may represent a TIC transcriptional state (Fig. 4B, Fig S3A).

To determine whether established TIC markers CD44^+^/CD24^low/-^ and ALDH1A1 could be used to confirm the identification of TIC transcriptional states in our data, we generated feature plots based on their expression (Fig. S3C). In all three xenografts, CD44 had no distinct expression pattern and CD24 was highly expressed in all cells in BCM-4272 and BCM-15006, but not expressed in many SUM159 cells. Furthermore, ALDH1A1 was not a distinct marker of any clusters.

To gain insights into the biological states of each cluster, we used the differentially expressed genes that define the cells in a given cluster relative to all other cells to perform GSEA (Supplementary Tables 4–6). SUM159-C7, 4272-C7, and 15006-C6, which we hypothesized may represent a TIC transcriptional state, were defined by enrichment of IFNγ response, IFNα response, and TNF-alpha signaling via NF-kB (Fig. 4D, Supplementary Table 3). To investigate the extent to which candidate TIC clusters share transcriptional features, we intersected the differentially expressed genes from each cluster and performed Fisher’s exact test for significance (Fig. 4E). SUM159-C7 and 4272-C7 shared 48 features (p=1.8e^−24^), SUM159-C7 and 15006-C6 shared 38 features (p=5.7e^−27^), and 4272-C7 and 15006-C6 shared 79 features (p=1.1e^−110^). In this process, we identified only 25 common up-regulated genes across all three clusters, that may represent a new TIC signature. A list of genes from all intersections is shown in Supplementary Table 7.

To determine whether these 25 genes participate in shared biological processes, we performed a functional protein-protein interaction analysis using the STRING database (53) (Fig. 4F). We found all of our genes, except GNB4, exist within the same protein-protein interaction network (PPI enrichment: p<1.0e^−16^), which suggests the genes in our TIC signature may be part of a biological process responsible their regulation. To uncover this biological process, we performed an over-representation analysis using the 25-gene TIC signature, which confirmed these genes are overrepresented in gene sets and pathways related to IFN signaling (Fig. 4F, Fig. S4).

To determine whether our 25-gene signature can effectively distinguish cell states within each xenograft, we used marker enrichment modeling (35) to generate similarity scores based on the expression of each gene in the 25-gene signature and visualized the results with a heatmap (Fig. 4G). In the PDX models, 4272-C7 and 15006-C6 clustered separately from every other cell state in their respective xenografts, demonstrating the 25-gene signature is unique to these clusters. In contrast, SUM159-C7 is a member of the same clade as cluster 5, indicating cluster 5 also is defined by the expression of IFN genes. To distinguish these two clusters, we performed differential gene expression analysis and found SUM159-C7 differentially expressed genes associated with TIC, including *TP63*, *POU5F1*, *ABCA5*, *FOXO1*, and *TCF4*, supporting our hypothesis that SUM159-C7 represents TIC (Fig. S3D).

To determine whether additional PDX models have a similar transcriptional state, we analyzed an independent dataset of three PDX models sequenced using isolated nuclei rather than whole cells and gave each cell a TIC score based on the expression of the genes in our 25-gene signature (Fig. S5). All three PDX models (BCM-4013, BCM-15166, and WHIM30) had a single cluster with a high TIC score relative to all other clusters. Thus, our TIC signature defines distinct cell states in three additional PDXs, suggesting this cell state may be represented in a wider cohort of TNBC PDXs.

To confirm that IFNs can induce EGFP in cells transduced with the STAT reporter, we treated SUM159 STAT LT, BCM-4272 4M67-EGFP, and BCM-15006 4M67-EGFP cells with IFNα or IFNγ and confirmed that IFNγ activated the reporter in all three models more robustly than did IFNα. IFNα did not activate the reporter in SUM159 (Fig. S6A). To determine whether IFN treatment activates genes that define the IFN-associated TIC transcriptional state, we input our 25 genes into INTERFEROME (54), an online database of IFN-regulated genes, which revealed all 25 genes are regulated by IFN (Fig. S6B).

To demonstrate these genes can be regulated by IFN in our models, we compared the expression of three upstream regulators of IFN signaling (PARP9, PARP14, STAT1) and three genes activated in response to IFN signaling (ADAR, OAS2, and BST2) from our 25 intersected TIC genes after IFN treatment to untreated controls by qRT-PCR (Fig. S6C). IFNα or IFNγ treatment increased the expression of all six genes relative to the untreated controls in all three TNBC models.

To evaluate whether any candidate TIC genes we identified have been previously reported, we compared intersections from any two xenograft models to five previously published TIC signatures that were derived using bulk RNA sequencing, including our own (30–34), and a previously published panel of stem cell and epithelial-to-mesenchymal transition markers (55) (Supplementary Table 8). None of the 25 genes we identified overlapped significantly with any of the six previously published TIC signatures. The Pece et al. (33) signature contained one of our TIC genes. LY6E, from the 4272-C7 and 15006-C6 intersection, whereas the Colacino et al. (55) custom panel only contained CDH3 (SUM159-C7 and 4272-C7 intersection) and MCL1 and ITGA6 (SUM159-C7 and 15006-C6 intersection). Furthermore, a comparison of these previously published TIC signatures did not reveal any significant gene-level commonalities, suggesting that single cell methods may be required for accurate identification of TIC.

### Interferon response genes are differentially expressed in a subset of epithelial cells in human breast cancer patient samples

To determine whether a similar IFN/STAT1 transcriptional state could be detected in human clinical samples, we analyzed scRNAseq data retrieved from a published human breast cancer cell atlas that contained TNBC, HER2^+^, ER^+^, pre-neoplastic (BRCA1^+/-^), and normal breast samples (37). To identify epithelial cells from breast cancer patients with a similar transcriptional state to our IFN/STAT1 TIC, we gave each cell a TIC score based on the expression of the genes in our 25-gene TIC signature (Fig. 5A–C). This identified cells in cluster 7 in TNBC patients (p=3.42e^−113^), cluster 6 in HER2^+^ patients (p=3.16e^−287^), and cluster 3 in ER^+^ patients (p=4.71e^−96^) had the highest TIC scores relative to cells in all other clusters. To determine whether established TIC markers CD44^+^/CD24^low/-^ and ALDH1A1 could be used to identify TIC transcriptional states in patient data, we generated feature plots based on their expression (Fig. S7A). In all three breast cancer subtypes, established TIC markers were not markers of any particular cluster, similar to our observations in PDX models.

**Figure 5:**
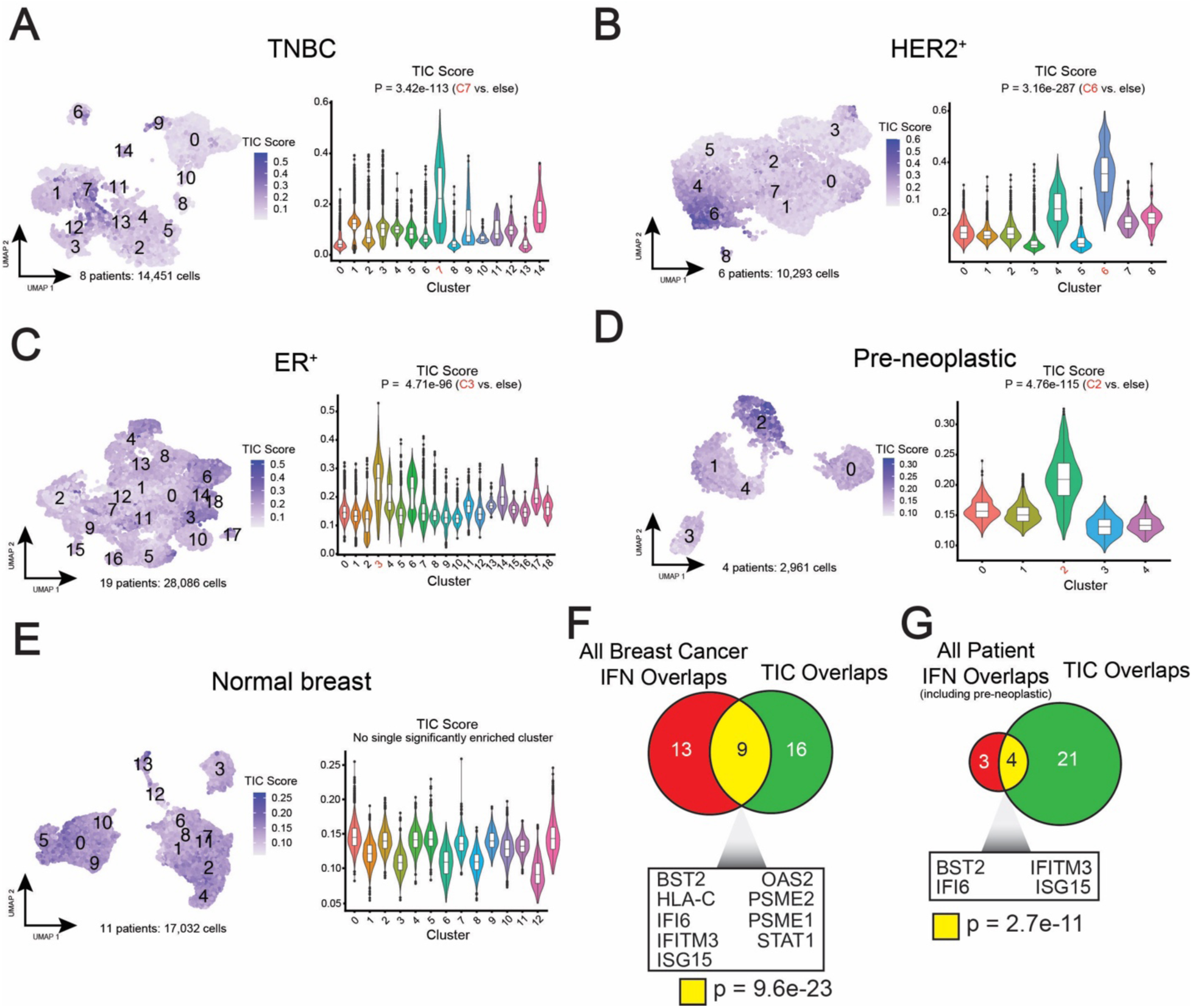
Interferon response genes are differentially expressed in a subset of epithelial cells in human breast cancer patient samples. A) UMAP visualization of eight human TNBC patient samples with feature plot and violin plot showing TIC score, derived from based on the expression of our 25-gene TIC signature. B) UMAP visualization of six human HER2^+^ patient samples with TIC score feature and violin plots. C) UMAP visualization of 19 human ER^+^ patient samples with TIC score feature and violin plots. D) UMAP visualization of four human pre-neoplastic (*BRCA1*^+/-^ tissue) patient samples with TIC score feature and violin plots. E) UMAP visualization of 11 human normal breast samples with TIC score feature and violin plots. In all TIC score violin plots, P values were calculated using a two-sided Wilcoxon test. F) The intersection between the common differentially expressed genes from clusters that were defined by a high TIC score across all three breast cancer subtypes and our 25-gene TIC signature. G) The intersection between the common differentially expressed genes from clusters that were defined by a high TIC score across all three breast cancer subtypes and pre-neoplastic tissue and our 25-gene TIC signature. The significance of gene list comparisons was calculated using Fisher’s exact test. IFN: interferon

To determine whether a similar transcriptional state exists in pre-neoplastic or normal breast tissue, we applied the same TIC scoring method (Fig. 5D–E). This revealed a similar IFN/STAT1 transcriptional state in pre-neoplastic tissue (cluster 2, p=4.76e^−115^), but not in normal breast. These results suggest the IFN/STAT1 transcriptional state is only present in breast cancer and pre-malignant tissue.

To determine the extent to which clusters with high TIC scores share transcriptional features with our 25-gene TIC signature, we intersected the genes from each cluster with a high TIC score from each breast cancer subtype, then intersected that gene list with our 25-genes TIC signature (Fig. 5F). This revealed the IFN/STAT1 transcriptional state from breast cancer patients shared 9 genes with our 25-gene signature (Fisher’s exact test: p=9.6e^−23^).

To determine whether this similarity extends to pre-neoplastic tissue, we intersected the genes from the high TIC score cluster from pre-neoplastic samples with the high TIC score intersected gene list from all breast cancer patients, then intersected that list with our 25-gene signature (Fig. 5G). This revealed the IFN/STAT1 transcriptional state from both breast cancer patients and pre-neoplastic tissue shared 4 genes with our 25-gene signature (Fisher’s exact test: p=2.7e^−11^).

To determine whether the IFN/STAT1 transcriptional state can be detected in individual breast cancer patients, we analyzed each breast cancer patient of each subtype with more than 1,500 non-cycling epithelial cells to increase our chances of identifying rare cell subpopulations, such as TIC (8) (Fig. S7B). We found that the IFN/STAT1 transcriptional state in most, but not all, patients analyzed, which is consistent with TIC heterogeneity. To establish the robustness of the TIC signature, we examined an additional cohort of TNBC patients from an additional human breast cancer scRNAseq dataset (38). We gave each cell a TIC score based on the expression of the genes in our 25-gene TIC signature (Fig. S8A), which revealed cluster 5 had a significantly higher TIC score relative to all other clusters (p=4.5e^−113^). Furthermore, the only patient with more than 1,500 non-cycling epithelial cells also had a cluster defined by a higher TIC score (cluster 4, p=2.3e^−03^) (Fig. S8B). Taken together, our findings reveal that the IFN/STAT1 signaling axis we observe in PDX models is present in human breast cancer patients.

### The IFN/STAT1 signaling axis is associated with worse overall survival and lack of response to chemotherapy in breast cancer patients

To evaluate the clinical significance of our TIC signature, we examine the association between high expression of our 25-gene TIC signature and overall survival using RNAseq datasets from KM-plotter (40). In all breast cancer patients, regardless of treatment status, the 25-gene TIC signature was associated with a worse overall survival (HR=1.46, p=0.0067) (Fig. 6A). This association with worse overall survival was more pronounced when stratifying for untreated patients (HR=2.33, p=0.0022), but was eliminated when stratifying for patients treated with chemotherapy (HR=0.77, p=0.23) (Fig. 6A), suggesting the TIC signature is only prognostic in untreated patients.

**Figure 6:**
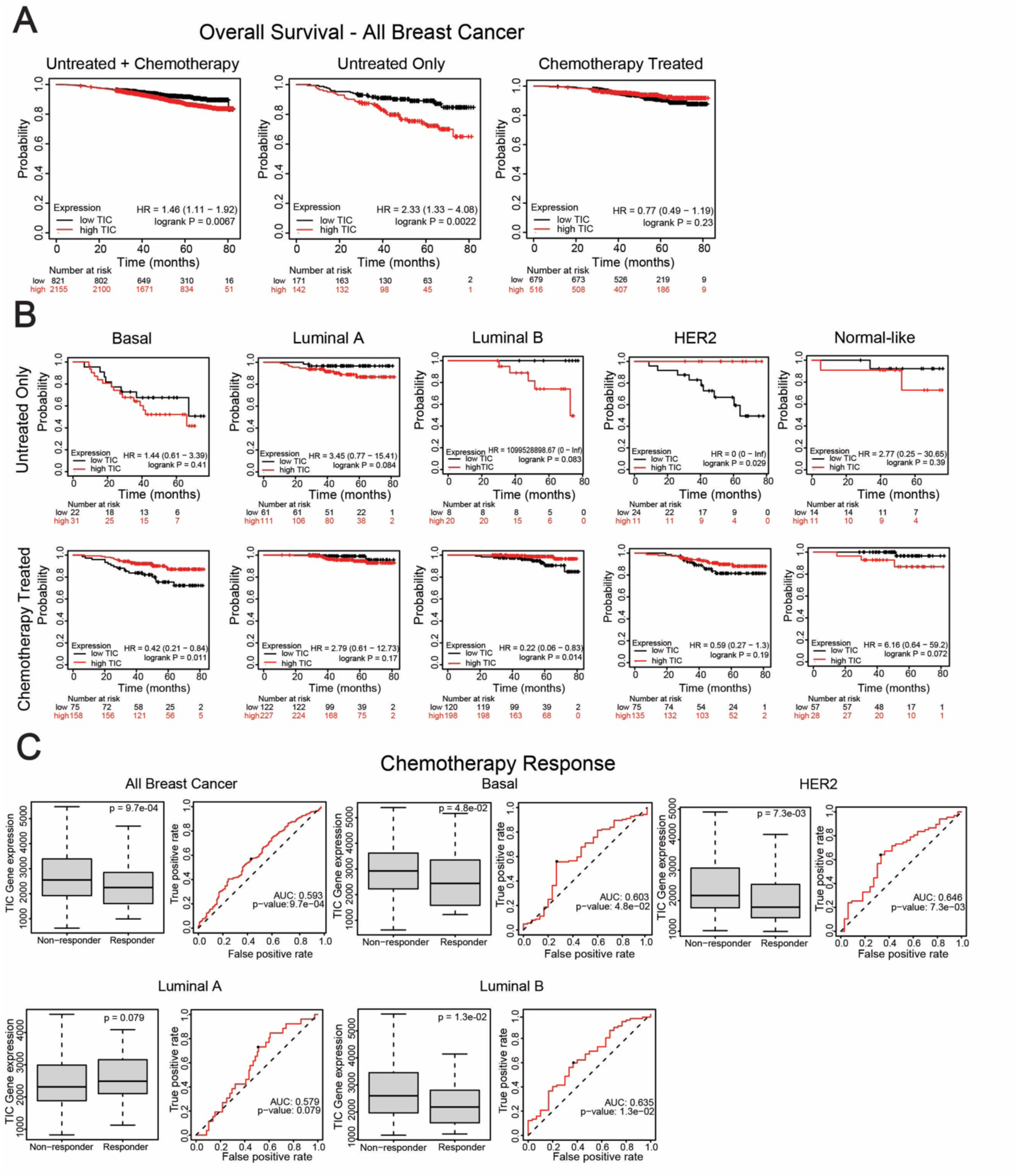
The IFN/STAT1 signaling axis is associated with worse overall survival and lack of response to chemotherapy in breast cancer patients. A) Kaplan-Meier plots showing overall survival between breast cancer patients with high or low expression of the 25-gene TIC signature including all patients (untreated + chemotherapy), only including untreated patients, and only including patients treated with chemotherapy. B) Kaplan-Meier plots showing overall survival between breast cancer patients with high or low expression of the 25-gene TIC signature comparing untreated patients and patients treated with chemotherapy by PAM50 subtype. C) ROC curve and box plots showing the association between the 25-gene TIC signature and chemotherapy response by breast cancer subtype.

Next, we stratified patients by PAM50 subtype and examined differences in overall survival in untreated patients compared to chemotherapy-treated patients (Fig. 6B). In basal, luminal A, and luminal B untreated patients, there was a trend of high TIC score with worse overall survival, which reduced in luminal A chemotherapy-treated patients or reversed in basal and luminal B chemotherapy-treated patients (Fig. 6B). High expression of TIC gene was associated with favorable overall survival in HER2 untreated patients, but this was reduced in chemotherapy-treated patients (Fig. 6B). Normal-like patients had a similar overall survival regardless of chemotherapy treatment (Fig. 6B).

We and others showed previously that TIC demonstrate intrinsic resistance to systemic chemotherapies. Thus, we evaluated whether expression of our 25-gene TIC signature is associated with response to chemotherapy in breast cancer patients using ROC plotter (41). Consistent with intrinsic chemotherapy resistance, high expression of our TIC signature was associated with non-responders, and was predictive of lack of response to chemotherapy (AUC: 0.593, p=9.7e^−04^) (Fig. 6C). Furthermore, this finding was consistent for all breast cancer subtypes, except for luminal A (Fig. 6C).

### BST2 allows TIC enrichment in breast cancer xenograft models

To demonstrate conclusively that the TIC transcriptional state identifies a subpopulation enriched for TIC function, we leveraged the observation that BST2 was a FACS-compatible cell surface found in the intersection between IFN/STAT1 transcriptional states from all breast cancer subtypes and pre-neoplastic tissue and our 25-gene signature. We performed analytical flow cytometry to examine the relationship between BST2 protein levels and TIC marker CD44^+^/CD24^low/-^ and ALDH activity. In SUM159 xenografts, BST2^+^ cells are ∼40% of the total tumor but represent 96% of CD44^+^/CD24^low/-^ cells and 58% of ALDH^+^ cells. In BCM-4272 xenografts, BST2^+^ cells represented ∼20% of the tumor, but 38% of CD44^+^/CD24^low/-^ cells and 92% of ALDH^+^ cells. These results indicate BST2 expression correlates with established TIC markers, suggesting BST2 may also allow enrichment for TIC.

To determine whether cells that highly express BST2 are enriched for mammosphere forming potential and/or self-renewal activity we dissociated SUM159 xenografts and FACS-enriched BST2^high^ (top 20%), BST2^low/-^ (bottom 20%), and mock sorted cells to test their primary and secondary mammosphere forming frequencies (Fig. 7B). BST2^high^ cells formed both primary and secondary mammospheres at a greater frequency than either the BST2^low/-^ cells or the mock sorted control, with no statistically significant difference between the mock control and BST2^low/-^.

**Figure 7:**
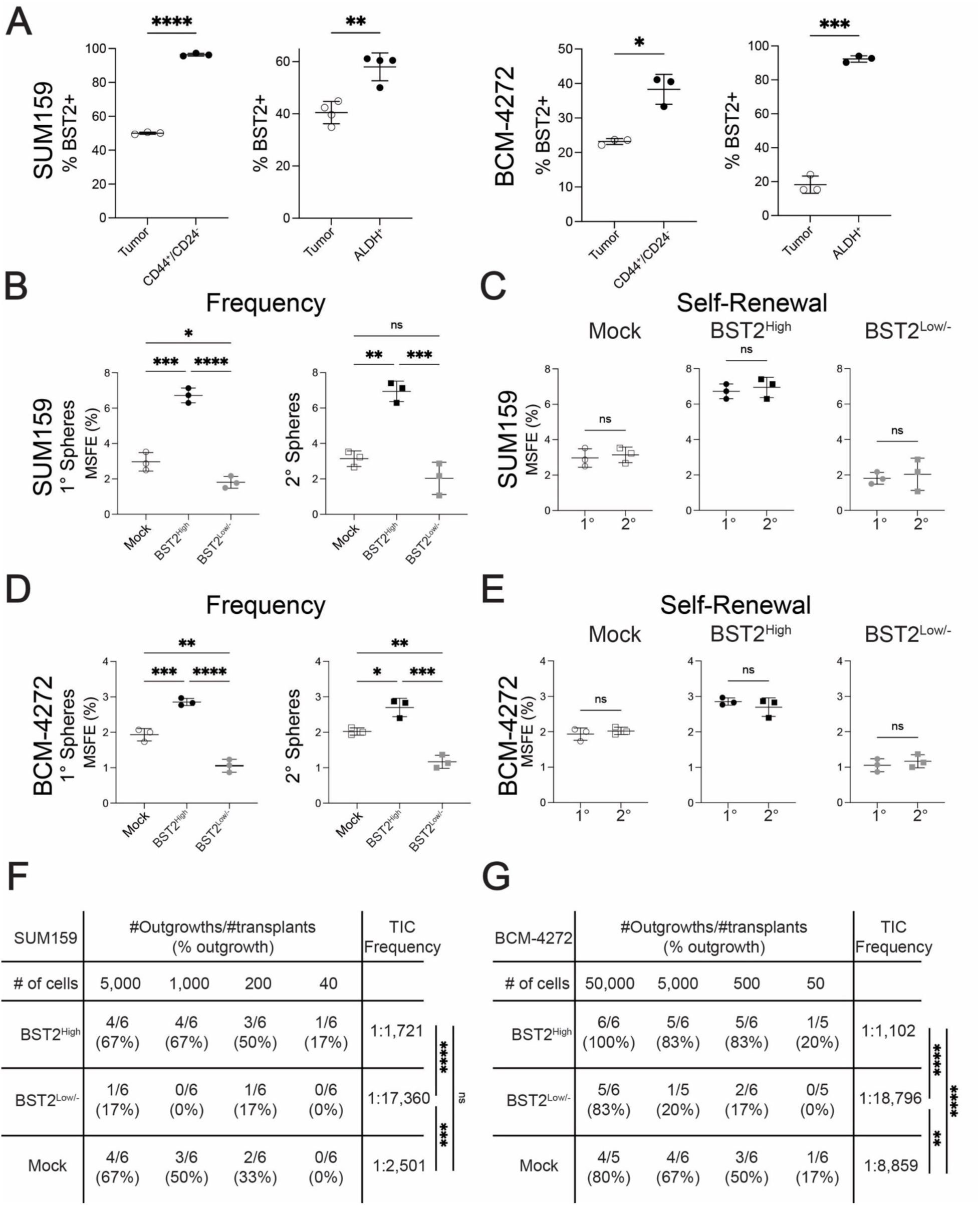
BST2 allows TIC enrichment in breast cancer xenograft models. A) Flow cytometry plot examining the correlation between BST2 and TIC markers CD44^+^/CD24^−^ and ALDEFLUOR activity in SUM159 and BCM-4272 xenografts. A two-tailed T-test was used to determine significance. B) SUM159 xenografts were dissociated, then the BST2 high, BST2 low, and mock-sorted populations were FACS enriched and grown as primary mammospheres (1°), then passaged and grown as secondary mammospheres (2°) (*n* = 3 biological replicates). One-way ANOVA and Tukey’s post hoc test were used to determine statistical significance. C) Differences in MSFE between primary and secondary mammospheres for each flow-sorted fraction. A two-tailed T-test was used to determine significance. D) BCM-4272 xenografts were dissociated, then the BST2 high, BST2 low, and mock-sorted populations were FACS enriched and grown as primary mammospheres, then passaged and grown as secondary mammospheres. MSFE of each cell population for each xenograft (*n* = 3 biological replicates). One-way ANOVA and Tukey’s post hoc test were used to determine statistical significance. E) Differences in MSFE between primary and secondary mammospheres for each flow-sorted fraction A two-tailed T-test was used to determine significance. F) Limiting dilution transplantation results of BST2 high, BST2 low, and mock-sorted populations from SUM159 xenografts. TIC frequency was calculated using extreme limiting dilution analysis and a Fisher’s exact test was used to determine statistical significance (25). G) Limiting dilution transplantation results of BST2 high, BST2 low, and mock-sorted populations from BCM-4272 xenografts. TIC frequency was calculated using extreme limiting dilution analysis and a Fisher’s exact test was used to determine statistical significance (25). *p < 0.05, **p < 0.01, ***p <0.001, ****p < 0.0001; values are mean ± SD.

To determine whether self-renewal capacity differed between the two populations, we compared the rate of MSFE between the primary and secondary mammospheres and found the BST2^high^, BST2^low/-^, and mock sorted control form mammospheres at the same rate over passaging (Fig. 7C) indicating no differences in self-renewal capacity. Similar results were observed for BCM-4272 (Fig. 7D and 7E). These results indicate that cells with high BST2 expression have enhanced MSFE, but not increased self-renewal capacity.

Finally, to test whether BST2 can be used to enrich TIC directly, we compared the tumor-initiating potential of BST2^high^ (top 20%), BST2^low/-^ (bottom 20%), and mock-sorted cells from SUM159 xenografts using a limiting-dilution transplantation assay (Fig. 7F). Notably, the BST2^high^ cells were the only population capable of forming tumors at all doses, with a TIC frequency of 1 in 1,721 cells. The BST2^low/-^ cells had a 10-fold lower TIC frequency of 1 in 17,370 cells, whereas the mock-sorted group had a modestly lower TIC frequency of 1 in 2,501 cells, likely due to the presence of TIC with lower BST2 expression present in the 60% of cells not assayed.

Qualitatively similar results were obtained using BCM-4272 xenografts using a limiting-dilution transplantation assay (Fig. 7G). The BST2^high^ cells were the only population capable of forming tumors at all doses, with a TIC frequency of 1 in 1,102 cells, a much higher TIC frequency than the mock sorted control, which had a TIC frequency of 1 in 8,859 cells. In BST2^low/-^ cells, tumor-initiating potential was depleted relative to the mock sorted control, with a TIC frequency of 1 in 18,796 cells. Taken together, our findings indicate BST2 can enrich for IFN/STAT1 signaling TIC.

### BST2 regulates TIC frequency in SUM159

To explore whether BST2 regulates TIC function, we generated two BST2 knockdown SUM159 cell lines (sh-BST2 #1 and sh-BST2 #2) (Fig. 8A) and compared their primary and secondary MSFE relative to a non-targeting control (Fig. 8B). BST2 knockdown reduced primary MSFE from 2.5% in the non-targeting control cells to 1.5% and 1.3% in shBST2 #1 and shBST2 #2, respectively, whereas BST2 knockdown reduced secondary MSFE from 3% in the non-targeting control cells to 1.6% and 1.2% in shBST2 #1 and shBST2 #2, respectively. These results demonstrate BST2 regulates mammosphere-forming frequency in SUM159 cells.

**Figure 8:**
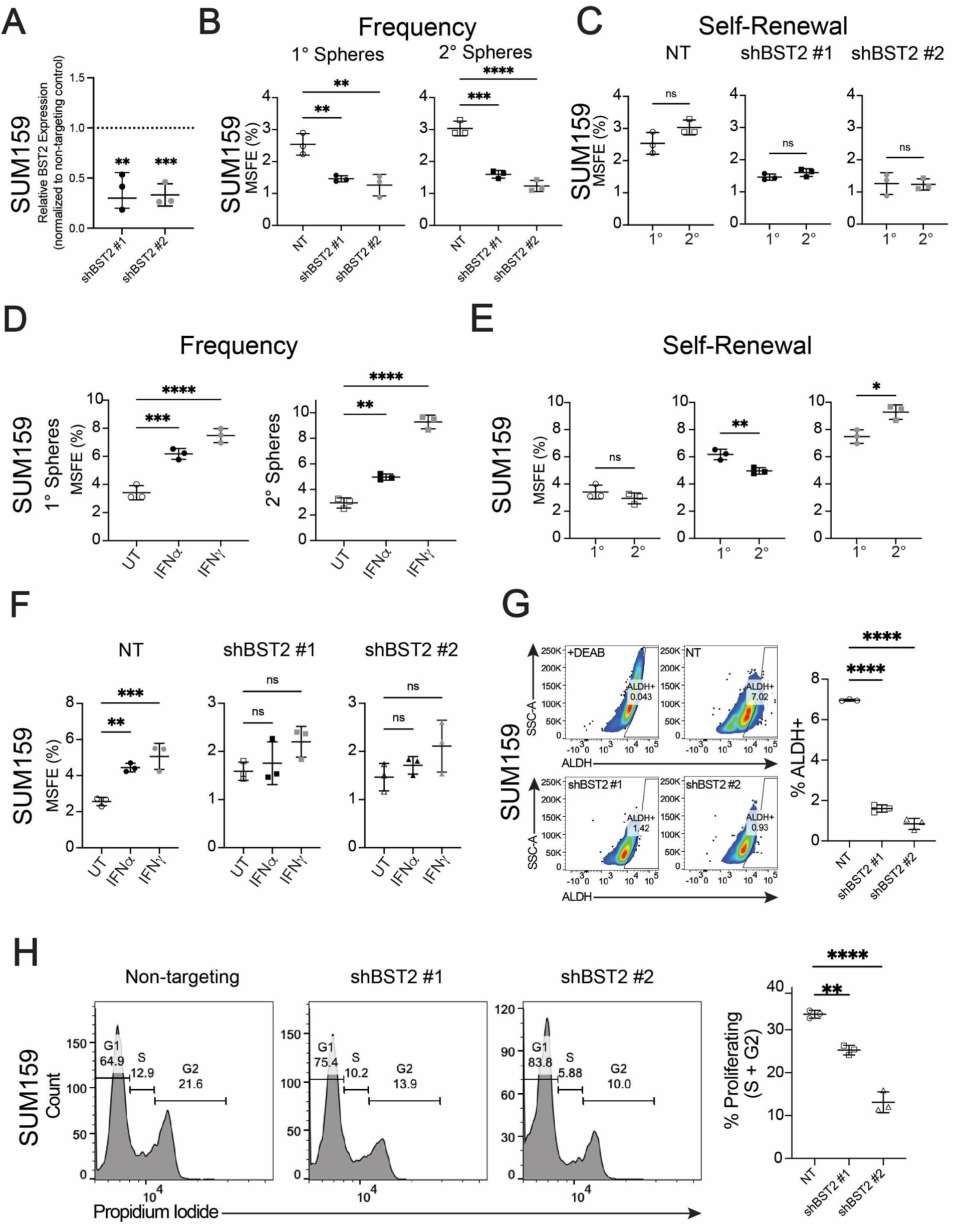
BST2 regulates TIC frequency in SUM159. A) BST2 was knocked down in SUM159 cells using two different shRNAs (shBST2 #1 and shBST2 #2), then BST2 knockdown was confirmed using Jess automated western blot. A two-tailed T-test was used to evaluate significance between each knockdown line and a non-targeting control (NT). B) SUM159 NT, shBST2 #1, and shBST2 #2 cells were grown as primary mammospheres (1°), then passaged and grown as secondary mammospheres (2°) (*n* = 3 biological replicates). One-way ANOVA and Tukey’s post hoc test were used to determine statistical significance. C) Differences in MSFE between primary and secondary mammospheres for each cell line. A two-tailed T-test was used to determine significance. D) SUM159 cells were grown as primary mammospheres (1°) in the presence of IFNα (100 ng/μL), IFNγ (100 ng/μL), or left untreated as a control, then passaged and grown as secondary mammospheres (2°) in the presence of IFNα (100 ng/μL), IFNγ (100 ng/μL), or left untreated as a control (*n* = 3 biological replicates). One-way ANOVA and Tukey’s post hoc test were used to determine statistical significance. E) Differences in MSFE between primary and secondary mammospheres for treatment. A two-tailed T-test was used to determine significance. F) SUM159 NT, shBST2 #1, and shBST2 #2 cells were grown as primary mammospheres (1°) in the presence of IFNα (100 ng/μL), IFNγ (100 ng/μL), or left untreated (UT) as a control (*n* = 3 biological replicates). One-way ANOVA and Tukey’s post hoc test were used to determine statistical significance. G) Representative flow cytometry plots illustrating ALDH activity in SUM159 NT, shBST2 #1, and shBST2 #2 cells and quantification of three biological replicates. H) Representative cell cycle flow cytometry plots using propidium iodide in SUM159 NT, shBST2 #1, and shBST2 #2 cells and quantification of three biological replicates. *p < 0.05, **p < 0.01, ***p <0.001, ****p < 0.0001; values are mean ± SD. 1°: primary mammospheres, 2°: secondary mammospheres, UT: untreated, IFN: interferon.

To determine whether BST2 regulates self-renewal, we compared the rate of MSFE between primary and secondary mammospheres in BST2 knockdown cells relative to a non-targeting control (Fig. 8C). We found both BST2 knockdown cells formed mammospheres at the same rate over passaging compared to the non-targeting control, indicating that BST2 does not regulate self-renewal. Taken together, our data suggest BST2 functionally regulates TIC frequency independent of self-renewal.

Because IFN regulates the expression of genes in our TIC signature, including BST2 (Fig. S6B– C), we grew SUM159 mammospheres in the presence of IFNα or IFNγ and compared primary and secondary mammosphere-forming frequency to an untreated control (Fig. 8D). Both IFNα and IFNγ increased primary mammosphere frequency to 6.2% and 7.5%, respectively, compared to the untreated control (3.4%). Similarly, IFNα and IFNγ increased secondary mammosphere frequency to 5% and 9.3%, respectively, compared to the untreated control (2.9%). However, only IFNγ increased rate of MSFE over passaging (from 7.5% to 9.3%), suggesting IFNγ regulates both TIC frequency and self-renewal in SUM159 (Fig. 8E).

To determine whether BST2 is responsible for the IFN-mediated increase in MSFE, we compared the effect of IFN on MSFE in two BST2 knockdown SUM159 cell lines (shBST2 #1 and shBST2 #2) compared to a non-targeting control (Fig. 8F). In the non-targeting control, IFNα increased MSFE to 4.4% and IFNγ increased MSFE to 5.1% relative to the untreated control (2.7%). In both BST2 knockdown lines, neither IFNα nor IFNγ increased MSFE relative to the untreated control, suggesting BST2 is a mediator of the IFN-mediated increase in MSFE.

ALDH activity is a TIC marker in SUM159 (56). To evaluate whether BST2 knockdown reduces ALDH activity, we performed an ALDEFLUOR assay in BST2 knockdown SUM159 cell lines relative to a non-targeting control. BST2 knockdown reduced ALDH activity from 7.0% in the non-targeting control cells to 1.6% and 0.8% in shBST2 #1 and shBST2 #2, respectively (Fig. 8G).

To determine whether this increase in TIC is due to proliferation, we compared cell cycle changes following BST2 knockdown. Of the non-targeting control cells, 33.7% were proliferative, whereas only 25.4% in shBST2 #1, and 13.1% in shBST2 #2, were proliferative, indicating BST2 knockdown reduces proliferation (Fig. 8H). Taken together, our data suggest that BST2 regulates TIC frequency in SUM159, perhaps through modulating proliferation.

## DISCUSSION

Herein, we extend our previous finding that STAT reporter activity enriches for TIC in two cell line xenograft models of claudin-low TNBC (11,57), and identified PDX BCM-4272 as an additional non-claudin-low model where STAT reporter activity is associated with TIC functions. Despite this finding, the reporter did not enrich cells with enhanced MSFE in the three other TNBC PDX models. Data are consistent with previous findings that STAT-mediated transcription has function(s) not necessarily associated with TIC. For example, STAT3 can promote immune evasion, metastasis, and chemoresistance (58), whereas STAT1 is associated with both tumor-promoting functions, such as immune suppression and metastasis, and tumor-suppressor functions, such as promoting apoptosis and immunosurveillance (59).

To address shortcomings of our original STAT-EGFP reporter, we developed the LT system to provide an inducible fluorescent label to tag STAT signaling TIC and other cells permanently as they undergo cell state changes. In addition, there are key questions in the cancer stem cell field that this system can address, including whether TIC from primary tumors function as metastasis-initiating cells (52,60), and whether enriched TIC populations following chemotherapy are the result of a therapeutic selective pressure, or are activated in response to the therapy (52,61).

We leveraged a synergy between scRNAseq, which has the power to detect rare cell populations, with our reporter systems to identify more precisely the TIC-enriched subpopulation of cells in multiple models of TNBC. The established TIC markers CD44^+^/ CD24^low/-^ and ALDH1A1 did not identify any unique cell states in our scRNAseq data, suggesting the expression of established TIC markers alone may not be sufficient to identify TIC in scRNAseq datasets. However, our analyses revealed that IFN response genes are differentially expressed in TIC in all three of our reporter-tagged TNBC, as well as three PDXs from an additional cohort, indicating the IFN/STAT1 signaling axis may be a common feature of TIC in a subset of TNBC. Critically, the IFN/STAT1 transcriptional state defines distinct cell states in all three breast cancer subtypes, as well as pre-neoplastic tissue. The absence of the IFN/STAT1 state in normal breast suggests this transcriptional state is a unique feature of some pre-neoplastic and neoplastic tissues, and may represent a candidate cancer-specific therapeutic target.

From an evolutionary perspective, the activation of IFN response genes in tumor cells presents multiple possibilities. On the one hand, TIC may have co-opted the IFN-mediated inflammatory response of the immune system to regulate TIC processes related to their frequency, such as differentiation, or cell state decisions (e.g. cellular quiescence/activation). Alternatively, and arguably more evolutionarily likely given that multicellular organisms arose from a single self-renewing cell, it is also possible that the immune system may have co-opted a stem cell pathway to mediate inflammatory responses.

Using the common genes across TIC states in all three PDX models in scRNAseq data, we derived a 25-gene TIC signature. Almost all genes we identified were not reported in previously published TIC signatures, all of which were derived using bulk RNA sequencing (30–34,55). Indeed, we found little agreement among the previously published TIC signatures themselves. This suggests that it is essential to combine TIC enrichment with scRNAseq to derive high-resolution TIC signatures.

There are limitations to consider when comparing bulk and single-cell samples. Bulk RNA sequencing is frequently performed at much higher coverage and depth than typical scRNAseq, which can allow detection of low expressing genes that may be missed by single-cell approaches. On the other hand, scRNAseq can dissect transcriptome heterogeneity, allowing discovery of rare cell types/states that would otherwise be obscured in bulk RNAseq.

With respect to translational potential, our 25-gene TIC signature was associated with worse overall survival in untreated breast cancer patients, but not in patients treated with chemotherapy. However, the signature was predictive of lack of response to chemotherapy, in line with previous findings that TIC are intrinsically resistant to systemic therapies (15). These data are also consistent with previous studies demonstrating enhanced expression of IFN genes following cytotoxic chemotherapy (62). Indeed, a tumor cell state defined by IFN gene expression was recently identified in TNBC scRNAseq, and was shown to be associated with tumors resistant to standard-of-care chemotherapies (63). This raises the possibility that TIC exploit the IFN/STAT1 axis to promote their activity in cancer, including mediating chemoresistance (64).

We identified BST2 as a candidate cell surface TIC marker, as it was differentially expressed in candidate TIC clusters across the three TNBC models we sequenced, and was a marker of clusters with a high TIC score in each breast cancer subtype, as well as pre-neoplastic tissue. BST2 is a transmembrane glycoprotein, whose expression is mediated by IFN, that is well-known for tethering newly-synthesized enveloped virions to the host cell plasma membrane (65) and activating NFκB to induce an antiviral state (66). To our knowledge, BST2 has never been associated with TIC in any tumor type.

Beyond utility as a TIC marker, our results demonstrate that BST2 is a regulator of TIC frequency in SUM159 and PDX BCM-4272, yet its knockdown in SUM159 did not affect self-renewal. This indicates BST2 must regulate TIC frequency by a different mechanism, such as quorum sensing. Quorum sensing, which is most commonly studied in bacteria, is a process of cell-to-cell communication that allows cells to share information about cell density and adjust their gene expression to regulate cell number (67). Organ-level quorum sensing has also been studied in stem cells, such as hair stem cell populations (68). To the extent that we have studied it, tumors appear to have specific TIC frequencies that are intrinsic properties of each tumor (69). If so, it stands to reason quorum sensing is one mechanism that regulates TIC frequency. As a cell surface protein, BST2 may be an integral component of the quorum sensing mechanism that regulates TIC frequency.

## ACKNOWLEDGMENTS

This work was supported, in part, by an F31 predoctoral fellowship (NCI CA278274) (to E.P.S), a U54 PDXNet PDX Development and Trials Center grant (NCI U54 CA224076) (to M.T.L.), an R01 grant (CA224867 to M.T.L. and H.L.F), a P30 Cancer Center Support Grant CA125123 (To C. Kent Osborne), a Core Facility Support Grant from the Cancer Research and Prevention Initiative of Texas RP170691 (to M.T.L.), the authors also acknowledge the joint participation by Diana Henry Helis Medical Research Foundation through its direct engagement in the continuous active conduct of medical research in conjunction with Baylor College of Medicine, the Komen for the Cure Promise grant, and a generous gift from the Korell family for the study of triple-negative breast cancer. S.S.Y. was supported by NIH grant R35GM133658 and Susan G. Komen Foundation grant CCR19609287. This project was also supported by the Cytometry and Cell Sorting Core at Baylor College of Medicine with funding from the CPRIT Core Facility Support Award (CPRIT-RP180672), the NIH (P30CA125123 and RR024574), the Single Cell Genomics Core at BCM with funding from NIH shared instrument grants (S10OD023469, S10OD025240) and P30EY002520, and the Genomic and RNA Profiling Core at Baylor College of Medicine with funding from NIH NCI (P30CA125123) and CPRIT (RP200504) grants. We thank Mu Wang for cloning the STAT lineage-tracing vectors as well as Dr. Koen Venken for helpful suggestions for vector design.

## Notes

### Competing Interest Statement

M.T.L is a Founder of, and an uncompensated Manager in, StemMed Holdings L.L.C., an uncompensated Limited Partner in StemMed Ltd. M.T.L is also a Founder of, and equity holder in, Tvardi Therapeutics. L.E.D. is a compensated employee of StemMed Ltd. Selected BCM PDX models described herein are exclusively licensed to StemMed Ltd. resulting in tangible property royalties to M.T.L. and L.E.D.

### Summary of Updates

The content is the same as previously. This revision is to fix a formatting issue

