## Supplemental Figures for "The interferon/STAT1 signaling axis is a common feature of tumor-initiating cells in breast cancer"

### SUPPLEMENTARY FIGURES

**Supplementary Table 1: pSTAT3 in breast PDX models**. Initial evaluation of 56 PDX models for pSTAT3 positivity by immunohistochemistry from tissue microarrays. A model was considered positive based on the detection of any pSTAT3-positive epithelial cells.

| PDX | pSTAT3 Status | Percent pSTAT3 Positive |
|---|---|---|
| BCM-3963 | Pos | 16.90% |
| BCM-3277 | Pos | 8.70% |
| BCM-7821 | Pos | 6.00% |
| BCM-3611 | Pos | 5.10% |
| BCM-7441 | Pos | 3.70% |
| BCM-5438 | Pos | 2.80% |
| BCM-3143 | Pos | 1.70% |
| BCM-3107 | Pos | 1.50% |
| BCM-0104 | Pos | 1.30% |
| HCI-028 | Pos | 1.3% |
| BCM-0132 | Pos | 1.20% |
| BCM-7649 | Pos | 1.00% |
| BCM-15046 | Pos | 1.00% |
| FCP-699 | Pos | 1.00% |
| MC1 | Pos | 0.80% |
| BCM-15008 | Pos | 0.70% |
| BCM-7482 | Pos | 0.60% |
| BCM-15115 | Pos | 0.60% |
| BCM-4013 | Pos | 0.50% |
| BCM-6257 | Pos | 0.50% |
| BCM-2147 | Pos | 0.50% |
| BCM-15030 | Pos | 0.50% |
| BCM-5998 | Pos | 0.30% |
| BCM-3469 | Pos | 0.27% |
| BCM-15057 | Pos | 0.25% |
| BCM-3561 | Pos | 0.20% |
| BCM-4272 | Pos | 0.20% |
| BCM-4913 | Pos | 0.20% |
| BCM-15051 | Pos | 0.15% |
| BCM-0002 | Pos | 0.14% |
| BCM-3204 | Pos | 0.13% |
| BCM-4849 | Pos | 0.09% |
| BCM-15006 | Pos | 0.09% |

| | | |
|---|---|---|
| BCM-3904 | Pos | 0.07% |
| BCM-15003 | Pos | 0.07% |
| BCM-3472 | Pos | 0.06% |
| BCM-4195 | Pos | 0.01% |
| HCI-002 | Pos | 0.01% |
| BCM-0046 | Neg | 0% |
| BCM-2665 | Neg | 0% |
| BCM-3887 | Neg | 0% |
| BCM-3936 | Neg | 0% |
| BCM-4175 | Neg | 0% |
| BCM-4664 | Neg | 0% |
| BCM-5471 | Neg | 0% |
| BCM-2277 | Neg | 0% |
| BCM-9161 | Neg | 0% |
| BCM-15029 | Neg | 0% |
| BCM-3104 | Neg | 0% |
| BCM-3613 | Neg | 0% |
| BCM-4169 | Neg | 0% |
| BCM-4888 | Neg | 0% |
| BCM-5097 | Neg | 0% |
| BCM-15034 | Neg | 0% |
| BCM-15100 | Neg | 0% |
| MDA-IBC3 | Neg | 0% |

**Supplementary Table 2: STAT reporter activity in breast PDX models**. Evaluation of 11 TNBC PDX models for STAT reporter activity (quantified from 5 representative images per model), including PDX biomarker expression and patient clinical features (retrieved from https://pdxportal.research.bcm.edu/pdxportal/?dswid=-3101).

| PDX Reporter Status | | | Patient Information | | |
|---|---|---|---|---|---|
| PDX | STAT Reporter | Mutation status | Metastasis Sites | Treatment Response Around Time of Sample Collection | Race/Ethnicity |
| BCM-4272 | 32.61% | TP53: frameshift variant<br>BRCA1: none<br>BRCA2: none | Bone, Liver, Ovary, Brain, Chest Wall, Lung | Partial Response (Docetaxel) | Hispanic or Latino |
| MC1 | 17.88% | TP53: missense variant<br>BRCA1: frameshift variant<br>BRCA2: none | No known metastasis | Unknown | White |
| BCM-7821 | 8.88% | TP53: frameshift variant<br>BRCA1: none<br>BRCA2: none | Pleura, Chest Wall, Lung | Partial Response (Docetaxel) | Black or African American |
| BCM-15006 | 3.93% | TP53: nonsense mutation<br>BRCA1: none<br>BRCA2: none | Bone | Partial Response (Doxorubicin, Cyclophosphamide)<br><br>Progressive disease (Paclitaxel, Carboplatin) | White |
| BCM-4013 | 1.19% | TP53: splice donor variant<br>BRCA1: none<br>BRCA2: missense variant | Chest Wall | Progressive Disease (Docetaxel) | Black or African American |
| BCM-3611 | 1.03% | TP53: nonsense mutation<br>BRCA1: missense variant<br>BRCA2: missense variant | No Known Metastasis | Progressive disease (Doxorubicin, Cyclophosphamide) | Black or African American |
| BCM-5097 | 0.68% | TP53: none<br>BRCA1: none<br>BRCA2: frameshift variant | Bone, Lung | Partial Response (Docetaxel) | White |
| HCI-028 | 0.47% | TP53: nonsense mutation<br>BRCA1: none<br>BRCA2: none | Bone, Lung, Ovary, Liver, Brain, Pleural Effusion | Progressive Disease (Adriamycin/Cyclophosphamide, Paclitaxel) | White |
| HCI-002 | 0.36% | TP53: none<br>BRCA1: none | Lymph Node | Unknown | White |

| | | | | | |
|---|---|---|---|---|---|
| | | BRCA2: none | | | |
| BCM-0046 | 0% | TP53: none<br>BRCA1: none<br>BRCA2: none | No Known Metastasis | Stable Disease (Docetaxel) | Hispanic or Latino |
| WHIM12 | 0% | TP53: missense variant<br>BRCA1: none<br>BRCA2: none | Pleural cavity, pericardium | Relapsed disease (5FU, Epirubicin, Cyclophosphamide, followed by Docetaxel) | White |

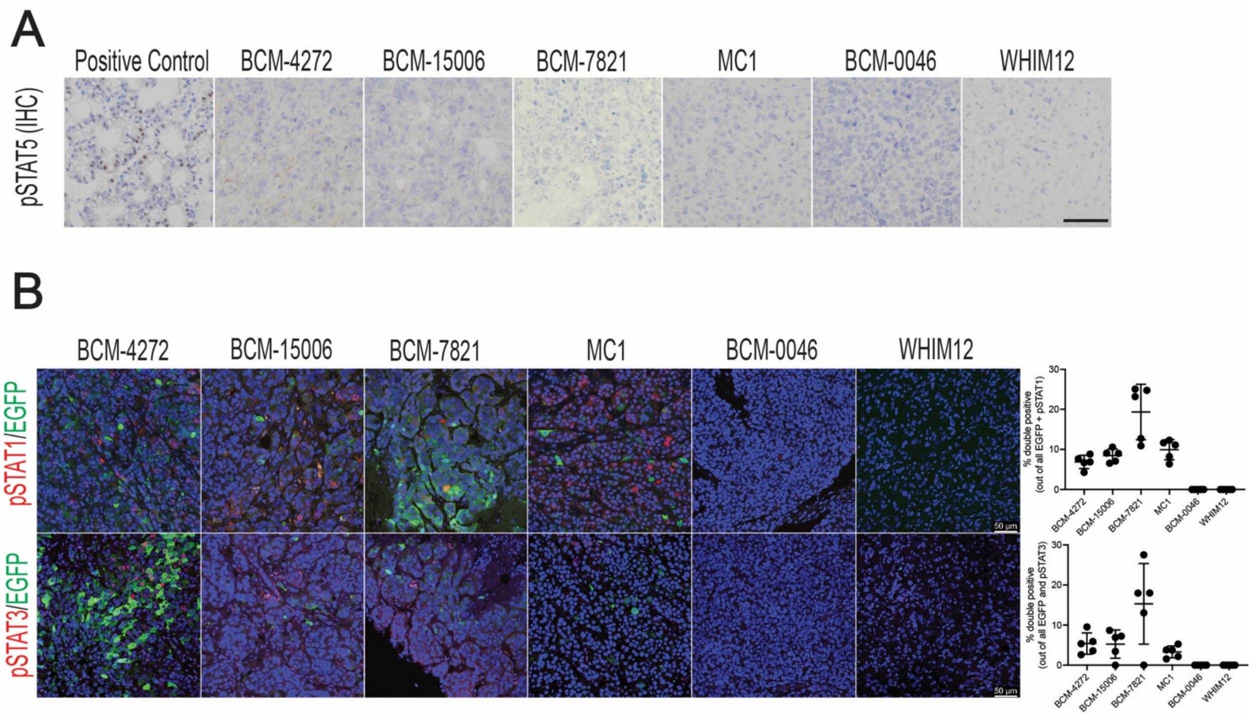

**Supplementary Figure 1: STAT reporter activity in breast PDX models.** A) Representative immunohistochemistry images showing levels of pSTAT5 in a panel of TNBC PDX models. A lactating mouse mammary gland was used as a positive control. B) Co-immunofluorescence staining for pSTAT1/EGFP and pSTAT3/EGFP with quantification revealing the extent of overlap between reporter positive cells and pSTAT1/3 levels.

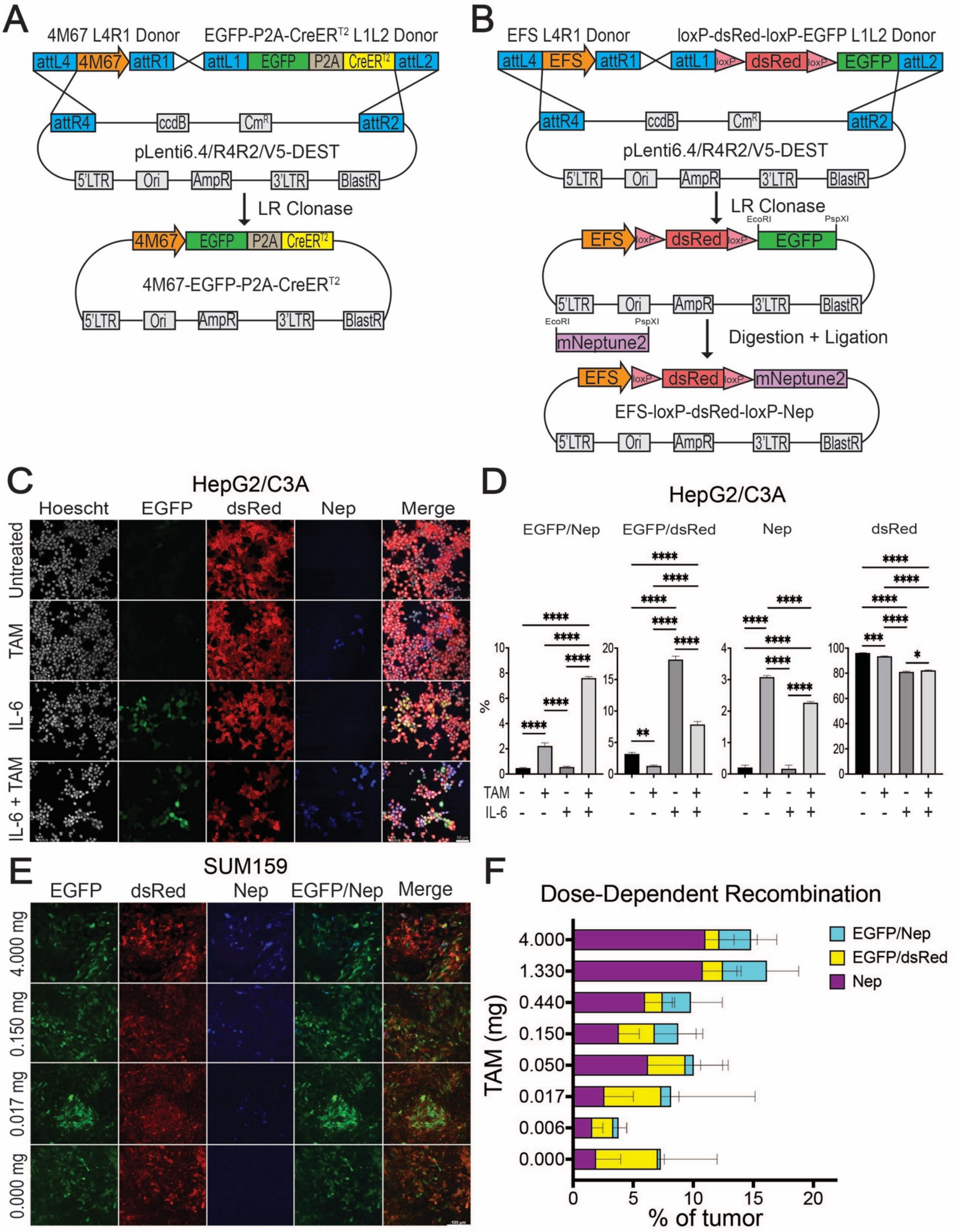
A
4M67 L4R1 Donor
EGFP-P2A-CreER$^{T2}$ L1L2 Donor
attL4
4M67
attR1
attL1
EGFP
P2A
CreER$^{T2}$
attL2
attR4
ccdB
Cm$^{R}$
attR2
pLenti6.4/R4R2/V5-DEST
5'LTR
Ori
AmpR
3'LTR
BlastR
LR Clonase
4M67-EGFP-P2A-CreER$^{T2}$
B
EFS L4R1 Donor
loxP-dsRed-loxP-EGFP L1L2 Donor
EFS
loxP
dsRed
EcoRI
PspXI
mNeptune2
Digestion + Ligation
EFS-loxP-dsRed-loxP-Nep
C
HepG2/C3A
Hoescht
EGFP
dsRed
Nep
Merge
Untreated
TAM
IL-6
IL-6 + TAM
D
HepG2/C3A
EGFP/Nep
EGFP/dsRed
Nep
dsRed
%
TAM
IL-6
E
SUM159
EGFP
dsRed
Nep
EGFP/Nep
Merge
4.000 mg
0.150 mg
0.017 mg
0.000 mg
F
Dose-Dependent Recombination
EGFP/Nep
EGFP/dsRed
Nep
TAM (mg)
4.000
1.330
0.440
0.150
0.050
0.017
0.006
0.000
% of tumor

**Supplementary Figure 2: STAT lineage tracing reporter provides a long-term fluorescent label to STAT signaling cells.** A) Generation of the 4M67-EGFP-P2A-CreERT2 reporter by LR Gateway® cloning. B) Generation of the EFS-loxP-dsRed-loxP-EGFP reporter by LR Gateway® cloning. Then, restriction cloning was used to insert mNeptune2 generating the EFS-loxP-dsRed-loxP-Nep dual color-switching reporter. C) In vitro validation of the STAT LT system in HepG2/C3A cells and D) quantification of each LT population by flow cytometry (*n* = 3). One-way ANOVA and Tukey's post hoc test were used to determine statistical significance. E) TAM dose-response curve illustrating dose-dependent activation of mNeptune2 expression. Images were taken two days after TAM treatment. F) Flow cytometry quantification of recombination illustrating the percentage of $EGFP^+/Nep^+$, $EGFP^+/dsRed^+$, and $mNeptune2^+$ cells in SUM159 xenografts (*n* = 5–8) two days after TAM treatment. One-way ANOVA and Tukey's post hoc test were used to determine statistical significance. *p < 0.05, ***p < 0.001, ****p < 0.0001; values are mean ± SD.

Nep: mNeptune2.

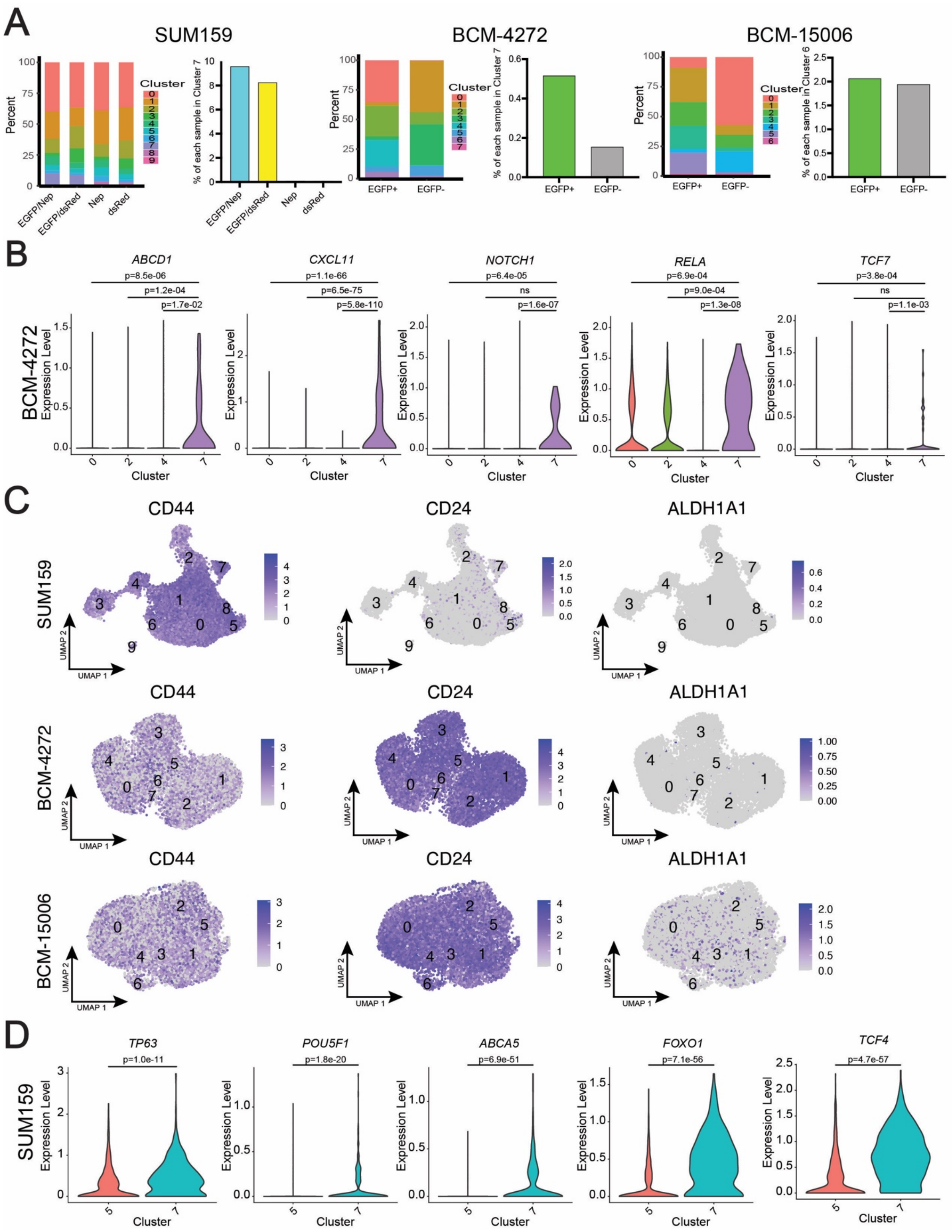

**Supplementary Figure 3: Interferon response genes are differentially expressed in TIC.** A) Feature plots for the expression of breast TIC markers CD44, CD24, and ALHH1A1 in SUM159

STAT LT, BCM-4272 4M67-EGFP, and BCM-15006 4M67-EGFP xenografts. B) Bar charts illustrating the distribution of each transcriptional state in each reporter population in SUM159, BCM-4272, and BCM-15006 xenografts, as well as a bar chart of the percentage TIC clusters represent within each reporter population. C) Violin plots illustrating the expression of genes associated with cancer stem cells that were differentially expressed in BCM-4272 cluster 7 relative to the three other clusters that were overrepresented in the $EGFP^{+}$ cells. A two-sided Wilcoxon test was used to determine statistical significance. D) Violin plots illustrating the expression of genes associated with cancer stem cells that were differentially expressed between SUM159 cluster 7 and SUM159 cluster 5. A two-sided Wilcoxon test was used to determine statistical significance.

**Supplementary Table 3: Interferon response genes are differentially expressed in TIC.** Reactome and GO Biological Processes GSEA of candidate TIC clusters for each xenograft model.

| | | Significantly Enriched Gene Sets from Differentially Expressed Up Genes | | | Significantly Enriched Gene Sets from Differentially Expressed Down Genes | | |
|---|---|---|---|---|---|---|---|
| | | Gene Set | p.adjust | qvalue | Gene Set | p.adjust | qvalue |
| SUM159 C7 | REACTOME | Interferon Signaling | 1.14E-11 | 9.41E-12 | KEAP1-NFE2L2 pathway | 7.13E-06 | 5.48E-06 |
| | | Interferon alpha/beta signaling | 4.02E-11 | 3.31E-11 | Diseases of signal transduction by growth factor receptors and secor | 1.06E-05 | 8.12E-06 |
| | | Interferon gamma signaling | 5.00E-09 | 4.13E-09 | Nuclear events mediated by NFE2L2 | 2.79E-05 | 2.14E-05 |
| | | Respiratory electron transport, ATP synthesis by | 5.55E-05 | 4.58E-05 | Attenuation phase | 2.79E-05 | 2.14E-05 |
| | | SARS-CoV Infections | 1.32E-04 | 1.09E-04 | Cellular response to heat stress | 2.79E-05 | 2.14E-05 |
| | GO_Biological_Process | Regulation of apoptotic signaling pathway | 2.38E-09 | 1.83E-09 | Regulation of mRNA processing | 3.91E-06 | 3.19E-06 |
| | | Regulation of peptidase activity | 4.08E-08 | 3.14E-08 | Establishment of protein localization to organelle | 3.91E-06 | 3.19E-06 |
| | | Negative regulation of apoptotic signaling pathw | 6.72E-08 | 5.16E-08 | Regulation of mRNA metabolic process | 2.43E-05 | 1.98E-05 |
| | | Regulation of endopeptidase activity | 6.82E-08 | 5.24E-08 | mRNA processing | 1.21E-04 | 9.87E-05 |
| | | Response to virus | 6.82E-08 | 5.24E-08 | RNA localization | 1.21E-04 | 9.87E-05 |
| BCM-4272 C7 | REACTOME | Interferon Signaling | 3.16E-33 | 2.41E-33 | Eukaryotic Translation Elongation | 2.04E-16 | 1.40E-16 |
| | | Interferon alpha/beta signaling | 7.28E-30 | 5.55E-30 | Peptide chain elongation | 3.35E-13 | 2.30E-13 |
| | | Interferon gamma signaling | 2.28E-19 | 1.74E-19 | Viral mRNA Translation | 3.35E-13 | 2.30E-13 |
| | | ER-Phagosome pathway | 6.68E-16 | 5.10E-16 | Selenocysteine synthesis | 3.35E-13 | 2.30E-13 |
| | | Antigen processing-Cross presentation | 1.88E-14 | 1.43E-14 | Eukaryotic Translation Termination | 3.35E-13 | 2.30E-13 |
| | GO_Biological_Process | Defense response to virus | 6.69E-35 | 5.68E-35 | Cytoplasmic translation | 1.22E-13 | 8.11E-11 |
| | | Defense response to symbiont | 6.69E-35 | 5.68E-35 | | | |
| | | Response to virus | 5.71E-34 | 4.85E-34 | | | |
| | | Negative regulation of viral process | 2.26E-23 | 1.92E-23 | | | |
| | | Regulation of viral process | 2.45E-23 | 2.08E-23 | | | |
| BCM-15006 C6 | REACTOME | Interferon Signaling | 1.15E-32 | 9.07E-33 | Eukaryotic Translation Elongation | 3.65E-52 | 2.81E-52 |
| | | Interferon alpha/beta signaling | 5.38E-30 | 4.26E-30 | Formation of a pool of free 40S subunits | 2.99E-46 | 2.30E-46 |
| | | Interferon gamma signaling | 8.93E-15 | 7.08E-15 | Peptide chain elongation | 9.91E-46 | 7.63E-46 |
| | | Antiviral mechanism by IFN-stimulated genes | 4.69E-09 | 3.72E-09 | L13a-mediated translational silencing of Ceruloplasmin expression | 3.02E-45 | 2.33E-45 |
| | | ISG15 antiviral mechanism | 6.45E-06 | 5.11E-06 | GTP hydrolysis and joining of the 60S ribosomal subunit | 3.21E-45 | 2.47E-45 |
| | GO_Biological_Process | Defense response to virus | 4.28E-34 | 3.55E-34 | Cytoplasmic translation | 6.89E-47 | 5.78E-47 |
| | | Defense response to symbiont | 4.28E-34 | 3.55E-34 | Ribosomal small subunit biogenesis | 1.05E-12 | 8.83E-13 |
| | | Response to virus | 6.01E-33 | 4.99E-33 | Ribosome biogenesis | 1.72E-12 | 1.44E-12 |
| | | Negative regulation of viral process | 2.22E-22 | 1.84E-22 | Ribonucleoprotein complex biogenesis | 1.96E-12 | 1.64E-12 |
| | | Regulation of viral process | 3.93E-22 | 3.26E-22 | rRNA processing | 5.22E-10 | 4.38E-10 |

**Supplementary Table 4: Interferon response genes are differentially expressed in TIC.**

Hallmark GSEA for every cluster from the SUM159 STAT LT xenograft.

| | Significantly Enriched Gene Sets from Differentially Expressed Up Genes | | | Significantly Enriched Gene Sets from Differentially Expressed Down Genes | | |
|---|---|---|---|---|---|---|
| | Gene Set | p.adjust | qvalue | Gene Set | p.adjust | qvalue |
| Cluster 0 | HALLMARK_MYC_TARGETS_V1 | 3.27E-07 | 2.37E-07 | HALLMARK_TNFA_SIGNALING_VIA_NFKB | 5.40E-21 | 4.18E-21 |
| | HALLMARK_REACTIVE_OXYGEN_SPECIES_PATHWAY | 1.98E-05 | 1.43E-05 | HALLMARK_UV_RESPONSE_DN | 3.34E-11 | 2.58E-11 |
| | HALLMARK_EPITHELIAL_MESENCHYMAL_TRANSITION | 2.69E-04 | 1.95E-04 | HALLMARK_EPITHELIAL_MESENCHYMAL_TRANSITION | 1.93E-08 | 1.49E-08 |
| | HALLMARK_INFLAMMATORY_RESPONSE | 8.80E-03 | 6.38E-03 | HALLMARK_HYPOXIA | 2.03E-07 | 1.57E-07 |
| | HALLMARK_OXIDATIVE_PHOSPHORYLATION | 8.80E-03 | 6.38E-03 | HALLMARK_TGF_BETA_SIGNALING | 2.11E-05 | 1.63E-05 |
| Cluster 1 | HALLMARK_UV_RESPONSE_DN | 1.02E-06 | 8.83E-07 | HALLMARK_OXIDATIVE_PHOSPHORYLATION | 3.68E-15 | 2.61E-15 |
| | HALLMARK_TNFA_SIGNALING_VIA_NFKB | 9.73E-05 | 8.40E-05 | HALLMARK_INTERFERON_ALPHA_RESPONSE | 3.10E-11 | 2.20E-11 |
| | HALLMARK_MYC_TARGETS_V2 | 1.83E-03 | 1.58E-03 | HALLMARK_INTERFERON_GAMMA_RESPONSE | 1.36E-08 | 9.62E-09 |
| | HALLMARK_G2M_CHECKPOINT | 2.36E-03 | 2.04E-03 | HALLMARK_EPITHELIAL_MESENCHYMAL_TRANSITION | 1.03E-05 | 7.29E-06 |
| | HALLMARK_MITOTIC_SPINDLE | 1.86E-02 | 1.60E-02 | HALLMARK_HYPOXIA | 1.03E-05 | 7.29E-06 |
| Cluster 2 | HALLMARK_TNFA_SIGNALING_VIA_NFKB | 7.42E-20 | 5.44E-20 | HALLMARK_MYC_TARGETS_V1 | 3.17E-20 | 2.32E-20 |
| | HALLMARK_INTERFERON_GAMMA_RESPONSE | 5.70E-14 | 4.17E-14 | HALLMARK_EPITHELIAL_MESENCHYMAL_TRANSITION | 1.15E-05 | 8.41E-06 |
| | HALLMARK_INTERFERON_ALPHA_RESPONSE | 3.12E-13 | 2.29E-13 | HALLMARK_TNFA_SIGNALING_VIA_NFKB | 1.80E-04 | 1.31E-04 |
| | HALLMARK_HYPOXIA | 5.34E-09 | 3.91E-09 | HALLMARK_MYC_TARGETS_V2 | 5.21E-04 | 3.80E-04 |
| | HALLMARK_APOPTOSIS | 3.10E-05 | 2.27E-05 | HALLMARK_APOPTOSIS | 4.28E-03 | 3.13E-03 |
| Cluster 3 | HALLMARK_EPITHELIAL_MESENCHYMAL_TRANSITION | 2.87.E-30 | 2.41E-30 | HALLMARK_TNFA_SIGNALING_VIA_NFKB | 3.98E-20 | 2.65E-20 |
| | HALLMARK_UV_RESPONSE_DN | 6.97.E-12 | 5.84E-12 | HALLMARK_P53_PATHWAY | 1.30E-07 | 8.69E-08 |
| | HALLMARK_TNFA_SIGNALING_VIA_NFKB | 1.09.E-03 | 9.11E-04 | HALLMARK_INFLAMMATORY_RESPONSE | 8.64E-07 | 5.75E-07 |
| | HALLMARK_ANGIOGENESIS | 5.63E-03 | 4.72E-03 | HALLMARK_INTERFERON_GAMMA_RESPONSE | 5.64E-06 | 3.76E-06 |
| | HALLMARK_ANDROGEN_RESPONSE | 2.84E-02 | 2.38E-02 | HALLMARK_APOPTOSIS | 2.87E-05 | 1.91E-05 |
| Cluster 4 | HALLMARK_MYC_TARGETS_V1 | 4.50.E-13 | 4.00E-13 | HALLMARK_INTERFERON_GAMMA_RESPONSE | 1.32E-21 | 9.64E-22 |
| | HALLMARK_MYC_TARGETS_V2 | 4.97.E-03 | 4.41E-03 | HALLMARK_TNFA_SIGNALING_VIA_NFKB | 8.34E-17 | 6.11E-17 |
| | HALLMARK_EPITHELIAL_MESENCHYMAL_TRANSITION | 1.39.E-02 | 1.23E-02 | HALLMARK_INTERFERON_ALPHA_RESPONSE | 3.50E-16 | 2.56E-16 |
| | HALLMARK_G2M_CHECKPOINT | 2.78E-02 | 2.47E-02 | HALLMARK_COMPLEMENT | 2.46E-07 | 1.80E-07 |
| | HALLMARK_ANDROGEN_RESPONSE | 3.04E-02 | 2.70E-02 | HALLMARK_HYPOXIA | 7.35E-07 | 5.38E-07 |
| Cluster 5 | HALLMARK_INTERFERON_GAMMA_RESPONSE | 2.27E-25 | 1.67E-25 | HALLMARK_EPITHELIAL_MESENCHYMAL_TRANSITION | 2.88E-10 | 2.17E-10 |
| | HALLMARK_INTERFERON_ALPHA_RESPONSE | 2.40E-18 | 1.76E-18 | HALLMARK_UV_RESPONSE_DN | 4.27E-09 | 3.21E-09 |
| | HALLMARK_TNFA_SIGNALING_VIA_NFKB | 9.96E-09 | 7.31E-09 | HALLMARK_TNFA_SIGNALING_VIA_NFKB | 5.32E-07 | 4.00E-07 |
| | HALLMARK_INFLAMMATORY_RESPONSE | 2.63E-06 | 1.93E-06 | HALLMARK_HYPOXIA | 1.67E-04 | 1.26E-04 |
| | HALLMARK_EPITHELIAL_MESENCHYMAL_TRANSITION | 8.08E-06 | 5.93E-06 | HALLMARK_APOPTOSIS | 2.61E-03 | 1.96E-03 |
| Cluster 6 | HALLMARK_HYPOXIA | 1.21E-24 | 9.33E-25 | HALLMARK_TNFA_SIGNALING_VIA_NFKB | 9.81E-20 | 6.95E-20 |
| | HALLMARK_TNFA_SIGNALING_VIA_NFKB | 3.77E-12 | 2.91E-12 | HALLMARK_MYC_TARGETS_V1 | 4.45E-08 | 3.16E-08 |
| | HALLMARK_MTORC1_SIGNALING | 5.36E-07 | 4.15E-07 | HALLMARK_INFLAMMATORY_RESPONSE | 3.37E-06 | 2.39E-06 |
| | HALLMARK_EPITHELIAL_MESENCHYMAL_TRANSITION | 5.64E-06 | 4.36E-06 | HALLMARK_EPITHELIAL_MESENCHYMAL_TRANSITION | 4.22E-04 | 2.99E-04 |
| | HALLMARK_APOPTOSIS | 1.19E-04 | 9.21E-05 | HALLMARK_MYC_TARGETS_V2 | 1.12E-02 | 7.91E-03 |
| Cluster 7 | HALLMARK_TNFA_SIGNALING_VIA_NFKB | 4.48E-23 | 3.54E-23 | HALLMARK_MYC_TARGETS_V1 | 5.47E-06 | 4.03E-06 |
| | HALLMARK_INTERFERON_GAMMA_RESPONSE | 3.00E-10 | 2.37E-10 | HALLMARK_UV_RESPONSE_DN | 3.75E-04 | 2.77E-04 |
| | HALLMARK_INTERFERON_ALPHA_RESPONSE | 6.92E-10 | 5.46E-10 | HALLMARK_EPITHELIAL_MESENCHYMAL_TRANSITION | 1.65E-03 | 1.22E-03 |
| | HALLMARK_APOPTOSIS | 5.94E-04 | 4.69E-04 | HALLMARK_HYPOXIA | 1.65E-03 | 1.22E-03 |
| | HALLMARK_INFLAMMATORY_RESPONSE | 1.32E-03 | 1.04E-03 | HALLMARK_MTORC1_SIGNALING | 7.08E-03 | 5.22E-03 |
| Cluster 8 | HALLMARK_EPITHELIAL_MESENCHYMAL_TRANSITION | 1.03E-08 | 7.30E-09 | HALLMARK_TNFA_SIGNALING_VIA_NFKB | 2.73E-09 | 2.18E-09 |
| | HALLMARK_TNFA_SIGNALING_VIA_NFKB | 7.42E-08 | 5.26E-08 | HALLMARK_MYC_TARGETS_V2 | 1.10E-08 | 8.79E-09 |
| | HALLMARK_INFLAMMATORY_RESPONSE | 1.37E-04 | 9.75E-05 | HALLMARK_MYC_TARGETS_V1 | 6.00E-07 | 4.80E-07 |
| | HALLMARK_APOPTOSIS | 1.37E-04 | 9.75E-05 | HALLMARK_G2M_CHECKPOINT | 4.03E-03 | 3.23E-03 |
| | HALLMARK_GLYCOLYSIS | 7.56E-04 | 5.36E-04 | HALLMARK_HYPOXIA | 4.03E-03 | 3.23E-03 |
| Cluster 9 | HALLMARK_OXIDATIVE_PHOSPHORYLATION | 3.10E-42 | 2.79E-42 | HALLMARK_MITOTIC_SPINDLE | 1.87E-10 | 1.38E-10 |
| | HALLMARK_MYC_TARGETS_V1 | 9.95E-16 | 8.94E-16 | HALLMARK_UV_RESPONSE_DN | 7.92E-10 | 5.84E-10 |
| | HALLMARK_DNA_REPAIR | 3.51E-06 | 3.15E-06 | HALLMARK_PROTEIN_SECRETION | 1.24E-09 | 9.16E-10 |
| | HALLMARK_REACTIVE_OXYGEN_SPECIES_PATHWAY | 1.24E-05 | 1.12E-05 | HALLMARK_TGF_BETA_SIGNALING | 6.65E-05 | 4.90E-05 |
| | HALLMARK_ADIPOGENESIS | 1.99E-02 | 1.79E-02 | HALLMARK_G2M_CHECKPOINT | 1.89E-03 | 1.40E-03 |

**Supplementary Table 5: Interferon response genes are differentially expressed in TIC.**

Hallmark GSEA for every cluster from the BCM-4272 4M67-EGFP xenograft.

| | Significantly Enriched Gene Sets from Differentially Expressed Up Genes | | | Significantly Enriched Gene Sets from Differentially Expressed Down Genes | | |
|---|---|---|---|---|---|---|
| | Gene Set | p.adjust | qvalue | Gene Set | p.adjust | qvalue |
| | HALLMARK_EPITHELIAL_MESENCHYMAL_TRANSITION | 2.8.E-11 | 2.21E-11 | HALLMARK_OXIDATIVE_PHOSPHORYLATION | 1.64E-21 | 1.07E-21 |
| | HALLMARK_TNFA_SIGNALING_VIA_NFKB | 6.2.E-07 | 4.93E-07 | HALLMARK_MYC_TARGETS_V1 | 6.73E-11 | 4.39E-11 |
| Cluster 0 | HALLMARK_INTERFERON_GAMMA_RESPONSE | 4.0.E-05 | 3.21E-05 | HALLMARK_E2F_TARGETS | 1.01E-06 | 6.59E-07 |
| | HALLMARK_HYPOXIA | 1.7.E-03 | 1.31E-03 | HALLMARK_MTORC1_SIGNALING | 7.51E-06 | 4.90E-06 |
| | HALLMARK_ANDROGEN_RESPONSE | 4.4.E-03 | 3.50E-03 | HALLMARK_MITOTIC_SPINDLE | 1.82E-05 | 1.19E-05 |
| | HALLMARK_OXIDATIVE_PHOSPHORYLATION | 1.53E-12 | 1.17E-12 | HALLMARK_TNFA_SIGNALING_VIA_NFKB | 7.79E-11 | 6.23E-11 |
| | HALLMARK_TNFA_SIGNALING_VIA_NFKB | 2.05E-06 | 1.58E-06 | HALLMARK_HYPOXIA | 5.01E-07 | 4.01E-07 |
| Cluster 1 | HALLMARK_CHOLESTEROL_HOMEOSTASIS | 2.83E-04 | 2.17E-04 | HALLMARK_UV_RESPONSE_DN | 5.94E-06 | 4.75E-06 |
| | HALLMARK_P53_PATHWAY | 2.10E-03 | 1.61E-03 | HALLMARK_INTERFERON_GAMMA_RESPONSE | 6.19E-06 | 4.95E-06 |
| | HALLMARK_HYPOXIA | 1.34E-02 | 1.03E-02 | HALLMARK_EPITHELIAL_MESENCHYMAL_TRANSITION | 1.12E-03 | 8.96E-04 |
| | HALLMARK_TNFA_SIGNALING_VIA_NFKB | 4.08E-14 | 2.70E-14 | HALLMARK_E2F_TARGETS | 5.54E-12 | 5.01E-12 |
| | HALLMARK_HYPOXIA | 2.65E-09 | 1.76E-09 | HALLMARK_G2M_CHECKPOINT | 2.42E-11 | 2.19E-11 |
| Cluster 2 | HALLMARK_INTERFERON_GAMMA_RESPONSE | 1.33E-04 | 8.81E-05 | HALLMARK_MITOTIC_SPINDLE | 4.65E-09 | 4.21E-09 |
| | HALLMARK_MTORC1_SIGNALING | 3.85E-04 | 2.56E-04 | HALLMARK_UV_RESPONSE_DN | 1.35E-05 | 1.22E-05 |
| | HALLMARK_ESTROGEN_RESPONSE_EARLY | 2.67E-03 | 1.77E-03 | | | |
| | HALLMARK_OXIDATIVE_PHOSPHORYLATION | 6.80E-07 | 6.07.E-07 | HALLMARK_TNFA_SIGNALING_VIA_NFKB | 1.20E-24 | 8.33E-25 |
| | HALLMARK_MYC_TARGETS_V1 | 6.20E-04 | 5.53.E-04 | HALLMARK_HYPOXIA | 3.13E-08 | 2.17E-08 |
| Cluster 3 | HALLMARK_REACTIVE_OXYGEN_SPECIES_PATHWAY | 3.17E-02 | 2.83.E-02 | HALLMARK_INTERFERON_GAMMA_RESPONSE | 5.16E-08 | 3.59E-08 |
| | | | | HALLMARK_INTERFERON_ALPHA_RESPONSE | 3.34E-05 | 2.32E-05 |
| | | | | HALLMARK_P53_PATHWAY | 4.23E-05 | 2.94E-05 |
| | HALLMARK_EPITHELIAL_MESENCHYMAL_TRANSITION | 6.48E-06 | 6.11.E-06 | HALLMARK_MTORC1_SIGNALING | 2.63E-22 | 1.77E-22 |
| | HALLMARK_TNFA_SIGNALING_VIA_NFKB | 3.56E-02 | 3.36.E-02 | HALLMARK_MYC_TARGETS_V1 | 2.55E-21 | 1.72E-21 |
| Cluster 4 | HALLMARK_UV_RESPONSE_DN | 3.91E-02 | 3.69.E-02 | HALLMARK_OXIDATIVE_PHOSPHORYLATION | 4.10E-13 | 2.76E-13 |
| | | | | HALLMARK_MYC_TARGETS_V2 | 6.21E-13 | 4.18E-13 |
| | | | | HALLMARK_UNFOLDED_PROTEIN_RESPONSE | 5.81E-07 | 3.91E-07 |
| | HALLMARK_MITOTIC_SPINDLE | 6.97E-04 | 5.72E-04 | HALLMARK_OXIDATIVE_PHOSPHORYLATION | 4.64E-14 | 3.42E-14 |
| | HALLMARK_APICAL_JUNCTION | 1.07E-03 | 8.77E-04 | HALLMARK_ADIPOGENESIS | 1.89E-08 | 1.39E-08 |
| Cluster 5 | HALLMARK_MYC_TARGETS_V1 | 9.45E-03 | 7.76E-03 | HALLMARK_INTERFERON_ALPHA_RESPONSE | 3.76E-06 | 2.77E-06 |
| | HALLMARK_ANDROGEN_RESPONSE | 9.45E-03 | 7.76E-03 | HALLMARK_INTERFERON_GAMMA_RESPONSE | 3.54E-04 | 2.61E-04 |
| | HALLMARK_CHOLESTEROL_HOMEOSTASIS | 4.06E-02 | 3.34E-02 | HALLMARK_TNFA_SIGNALING_VIA_NFKB | 3.54E-04 | 2.61E-04 |
| | HALLMARK_MYC_TARGETS_V1 | 1.29E-14 | 1.08E-14 | HALLMARK_TNFA_SIGNALING_VIA_NFKB | 1.59E-09 | 1.11E-09 |
| | HALLMARK_UNFOLDED_PROTEIN_RESPONSE | 1.38E-04 | 1.16E-04 | HALLMARK_UV_RESPONSE_DN | 1.96E-08 | 1.36E-08 |
| Cluster 6 | HALLMARK_OXIDATIVE_PHOSPHORYLATION | 3.99E-04 | 3.34E-04 | HALLMARK_EPITHELIAL_MESENCHYMAL_TRANSITION | 5.39E-06 | 3.74E-06 |
| | HALLMARK_MTORC1_SIGNALING | 2.84E-03 | 2.38E-03 | HALLMARK_ESTROGEN_RESPONSE_EARLY | 5.39E-06 | 3.74E-06 |
| | HALLMARK_MYC_TARGETS_V2 | 3.19E-03 | 2.67E-03 | HALLMARK_CHOLESTEROL_HOMEOSTASIS | 7.61E-05 | 5.29E-05 |
| | HALLMARK_INTERFERON_GAMMA_RESPONSE | 1.13E-56 | 9.22E-57 | HALLMARK_OXIDATIVE_PHOSPHORYLATION | 2.41E-09 | 1.77E-09 |
| | HALLMARK_INTERFERON_ALPHA_RESPONSE | 3.15E-52 | 2.56E-52 | HALLMARK_HYPOXIA | 1.00E-03 | 7.37E-04 |
| Cluster 7 | HALLMARK_TNFA_SIGNALING_VIA_NFKB | 9.79E-05 | 7.96E-05 | HALLMARK_ESTROGEN_RESPONSE_EARLY | 1.29E-03 | 9.52E-04 |
| | HALLMARK_ALLOGRAFT_REJECTION | 1.87E-02 | 1.52E-02 | HALLMARK_ADIPOGENESIS | 1.83E-03 | 1.35E-03 |
| | HALLMARK_COMPLEMENT | 3.13E-02 | 2.55E-02 | HALLMARK_TNFA_SIGNALING_VIA_NFKB | 2.72E-03 | 2.00E-03 |

**Supplementary Table 6: Interferon response genes are differentially expressed in TIC.**

Hallmark GSEA for every cluster from the BCM-15006 4M67-EGFP xenograft.

| | Significantly Enriched Gene Sets from Differentially Expressed Up Genes | | | Significantly Enriched Gene Sets from Differentially Expressed Down Genes | | |
|---|---|---|---|---|---|---|
| | Gene Set | p.adjust | qvalue | Gene Set | p.adjust | qvalue |
| Cluster 0 | HALLMARK_MYC_TARGETS_V1 | 1.33E-07 | 1.08E-07 | HALLMARK_MYC_TARGETS_V1 | 1.64E-21 | 1.21E-21 |
| | HALLMARK_MTORC1_SIGNALING | 1.13E-03 | 9.23E-04 | HALLMARK_MTORC1_SIGNALING | 7.78E-14 | 5.73E-14 |
| | HALLMARK_OXIDATIVE_PHOSPHORYLATION | 2.03E-02 | 1.66E-02 | HALLMARK_E2F_TARGETS | 1.64E-13 | 1.21E-13 |
| | HALLMARK_TNFA_SIGNALING_VIA_NFKB | 2.03E-02 | 1.66E-02 | HALLMARK_MYC_TARGETS_V2 | 3.00E-11 | 2.21E-11 |
| | HALLMARK_APOPTOSIS | 2.03E-02 | 1.66E-02 | HALLMARK_UNFOLDED_PROTEIN_RESPONSE | 3.22E-10 | 2.37E-10 |
| Cluster 1 | HALLMARK_OXIDATIVE_PHOSPHORYLATION | 5.56E-06 | 4.75E-06 | HALLMARK_MITOTIC_SPINDLE | 2.38E-07 | 1.86E-07 |
| | HALLMARK_MYC_TARGETS_V1 | 2.75E-04 | 2.35E-04 | HALLMARK_TNFA_SIGNALING_VIA_NFKB | 2.04E-06 | 1.59E-06 |
| | HALLMARK_APOPTOSIS | 2.41E-03 | 2.06E-03 | HALLMARK_UV_RESPONSE_DN | 1.51E-05 | 1.17E-05 |
| | HALLMARK_REACTIVE_OXYGEN_SPECIES_PATHWAY | 2.27E-02 | 1.94E-02 | HALLMARK_PROTEIN_SECRETION | 4.92E-04 | 3.83E-04 |
| | HALLMARK_HYPOXIA | 2.86E-02 | 2.44E-02 | HALLMARK_TGF_BETA_SIGNALING | 4.92E-04 | 3.83E-04 |
| Cluster 2 | HALLMARK_TNFA_SIGNALING_VIA_NFKB | 1.13E-05 | 8.93E-06 | HALLMARK_INTERFERON_ALPHA_RESPONSE | 4.81E-06 | 3.75E-06 |
| | HALLMARK_APOPTOSIS | 8.35E-05 | 6.59E-05 | HALLMARK_ADIPOGENESIS | 2.18E-04 | 1.70E-04 |
| | HALLMARK_EPITHELIAL_MESENCHYMAL_TRANSITION | 1.09E-02 | 8.59E-03 | HALLMARK_ESTROGEN_RESPONSE_LATE | 2.18E-04 | 1.70E-04 |
| | HALLMARK_IL2_STAT5_SIGNALING | 1.93E-02 | 1.52E-02 | HALLMARK_ESTROGEN_RESPONSE_EARLY | 2.50E-04 | 1.95E-04 |
| | HALLMARK_INTERFERON_ALPHA_RESPONSE | 1.93E-02 | 1.52E-02 | HALLMARK_OXIDATIVE_PHOSPHORYLATION | 2.50E-04 | 1.95E-04 |
| Cluster 3 | HALLMARK_TNFA_SIGNALING_VIA_NFKB | 9.73E-04 | 8.02E-04 | HALLMARK_MYC_TARGETS_V1 | 9.29E-09 | 6.06E-09 |
| | HALLMARK_APOPTOSIS | 1.69E-03 | 1.39E-03 | HALLMARK_OXIDATIVE_PHOSPHORYLATION | 7.00E-08 | 4.57E-08 |
| | HALLMARK_ANDROGEN_RESPONSE | 9.21E-03 | 7.59E-03 | HALLMARK_E2F_TARGETS | 2.57E-07 | 1.68E-07 |
| | HALLMARK_EPITHELIAL_MESENCHYMAL_TRANSITION | 9.21E-03 | 7.59E-03 | HALLMARK_MTORC1_SIGNALING | 3.95E-05 | 2.58E-05 |
| | HALLMARK_CHOLESTEROL_HOMEOSTASIS | 1.36E-02 | 1.12E-02 | HALLMARK_MYC_TARGETS_V2 | 2.20E-04 | 1.43E-04 |
| Cluster 4 | HALLMARK_MYC_TARGETS_V1 | 3.09E-15 | 2.32E-15 | HALLMARK_TNFA_SIGNALING_VIA_NFKB | 1.13E-15 | 7.85E-16 |
| | HALLMARK_E2F_TARGETS | 2.17E-03 | 1.63E-03 | HALLMARK_INTERFERON_ALPHA_RESPONSE | 9.20E-10 | 6.39E-10 |
| | HALLMARK_CHOLESTEROL_HOMEOSTASIS | 2.17E-03 | 1.63E-03 | HALLMARK_INTERFERON_GAMMA_RESPONSE | 9.20E-10 | 6.39E-10 |
| | HALLMARK_MTORC1_SIGNALING | 2.14E-02 | 1.61E-02 | HALLMARK_HYPOXIA | 3.13E-08 | 2.17E-08 |
| | HALLMARK_TNFA_SIGNALING_VIA_NFKB | 2.14E-02 | 1.61E-02 | HALLMARK_APOPTOSIS | 1.00E-06 | 6.97E-07 |
| Cluster 5 | HALLMARK_TNFA_SIGNALING_VIA_NFKB | 2.77E-10 | 2.11E-10 | HALLMARK_MITOTIC_SPINDLE | 4.74E-06 | 3.40E-06 |
| | HALLMARK_INTERFERON_GAMMA_RESPONSE | 6.40E-04 | 4.87E-04 | HALLMARK_ESTROGEN_RESPONSE_EARLY | 2.51E-04 | 1.80E-04 |
| | HALLMARK_APOPTOSIS | 1.12E-03 | 8.54E-04 | HALLMARK_G2M_CHECKPOINT | 3.23E-04 | 2.31E-04 |
| | HALLMARK_HYPOXIA | 3.45E-03 | 2.63E-03 | HALLMARK_ESTROGEN_RESPONSE_LATE | 8.43E-04 | 6.04E-04 |
| | HALLMARK_ANDROGEN_RESPONSE | 4.00E-03 | 3.04E-03 | HALLMARK_E2F_TARGETS | 1.22E-03 | 8.75E-04 |
| Cluster 6 | HALLMARK_INTERFERON_ALPHA_RESPONSE | 7.11E-46 | 5.76E-46 | HALLMARK_OXIDATIVE_PHOSPHORYLATION | 5.77E-14 | 4.73E-14 |
| | HALLMARK_INTERFERON_GAMMA_RESPONSE | 2.28E-41 | 1.85E-41 | HALLMARK_MYC_TARGETS_V1 | 2.75E-08 | 2.26E-08 |
| | HALLMARK_TNFA_SIGNALING_VIA_NFKB | 7.88E-05 | 6.38E-05 | HALLMARK_E2F_TARGETS | 1.35E-04 | 1.11E-04 |
| | HALLMARK_APOPTOSIS | 4.13E-02 | 3.34E-02 | HALLMARK_MYC_TARGETS_V2 | 3.39E-04 | 2.78E-04 |
| | | | | HALLMARK_DNA_REPAIR | 3.45E-04 | 2.83E-04 |

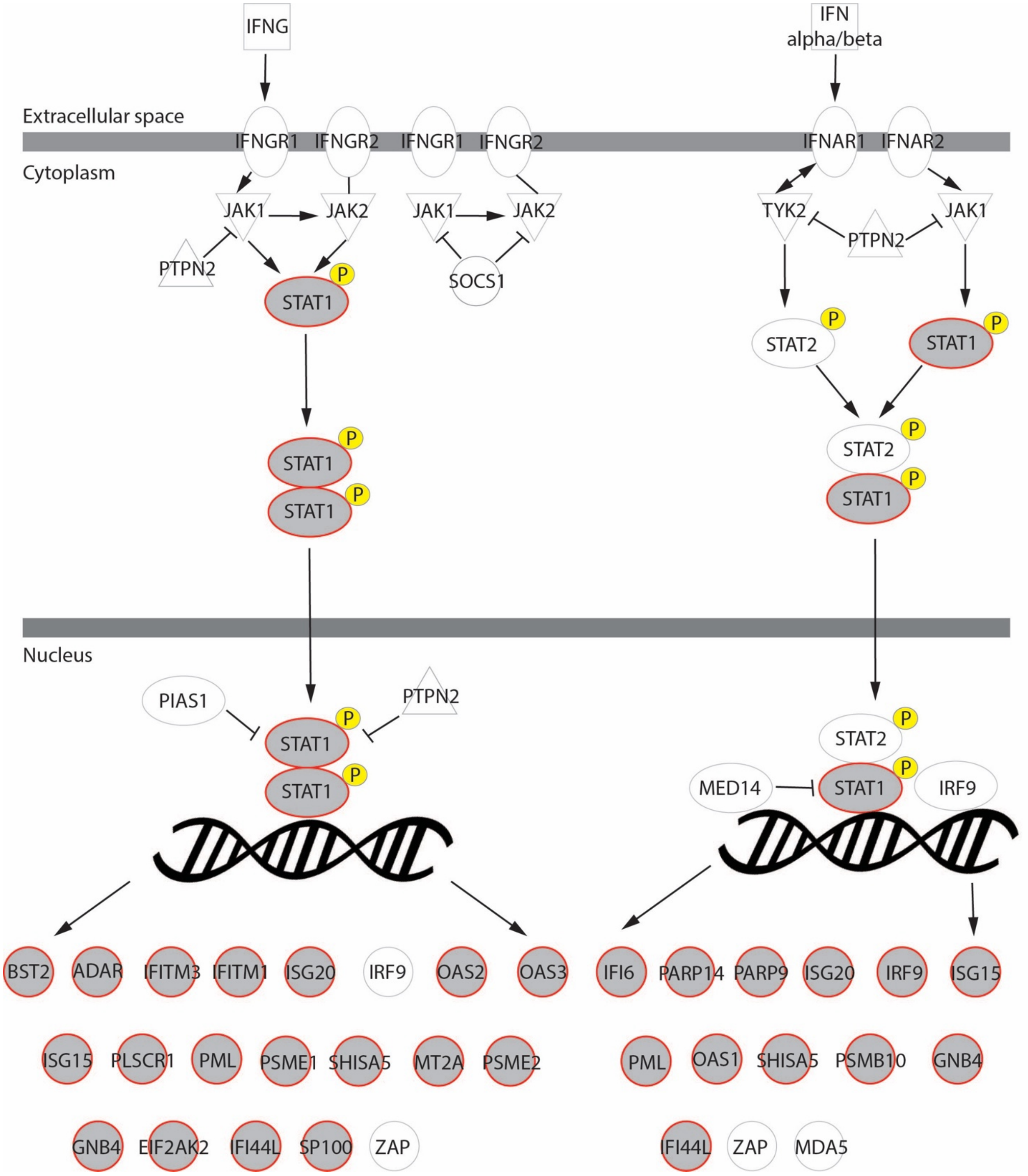

**Supplementary Figure 4: Interferon response genes are differentially expressed in TIC.** Interferon pathway map containing all 25 genes (except HLA-C) common across three candidate TIC clusters that participate in the interferon signaling pathway and/or are known outputs of the interferon signaling pathway. Grey shaded circles represent genes from our 25-gene list.

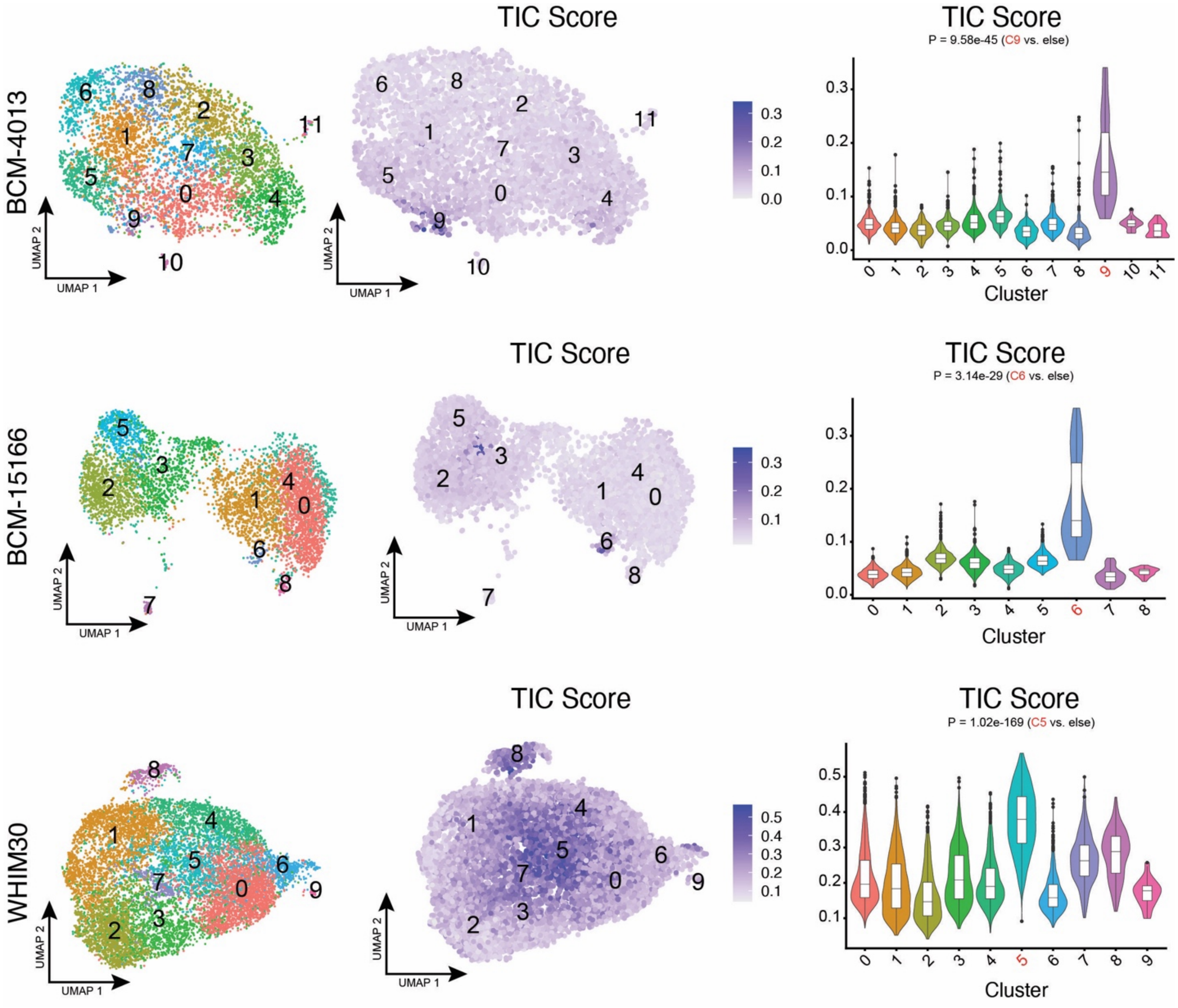

**Supplementary Figure 5: Interferon response genes are differentially expressed in TIC.** Analysis of an independent cohort of three breast PDX models. Each cell was given a TIC score based on the expression of the 25 genes common across TIC cell states in our three xenograft models. In all graphs, P values were calculated using a two-sided Wilcoxon test.

A

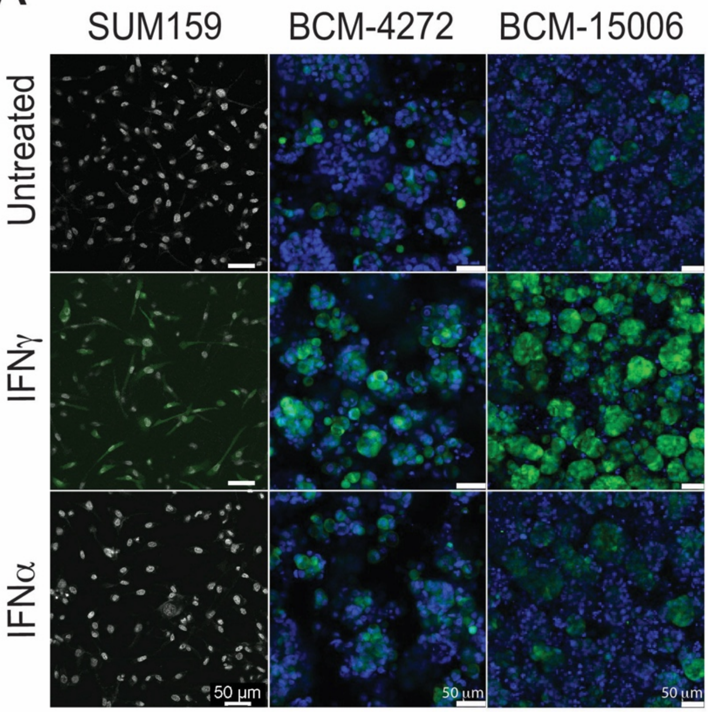

B

| INTERFEROME Homo sapiens statistics | | | | |
| --- | --- | --- | --- | --- |
| **Data base contents** | **Interferon Type I** | **Interferon Type II** | **Interferon Type III** | **Total** |
| Experiments | 40 | 29 | 2 | 60 |
| Datasets | 107 | 65 | 22 | 194 |
| Genes with p-value <0.05 and fold change > 2 | 7350 | 9768 | 184 | 12614 |
| **25-gene TIC signature** | 12 | | | 25 |
| | 13 | | | |

C

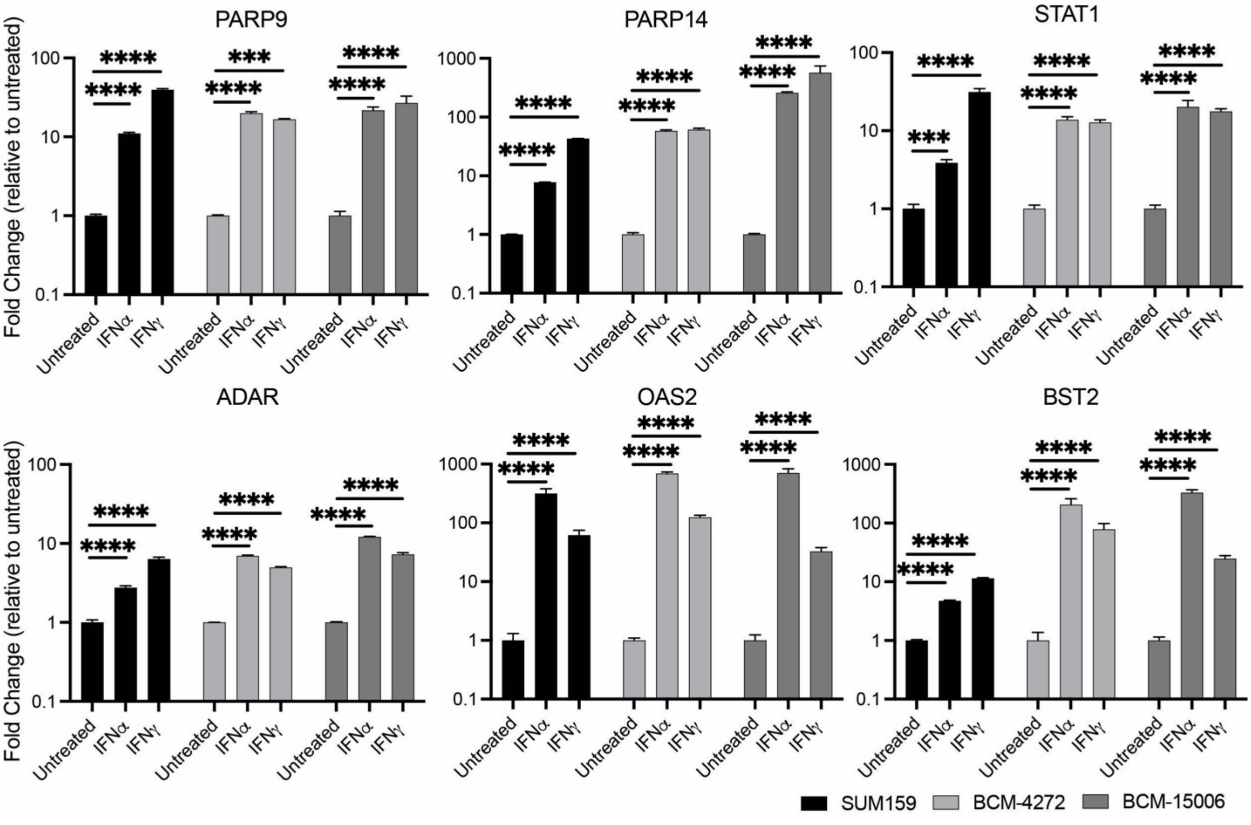

**Supplementary Figure 6: Interferon response genes are differentially expressed in TIC.** A) 4M67 reporter activity was evaluated by confocal microscopy in SUM159, BCM-4272, and BCM-15006 cells treated with interferon alpha (IFN$\alpha$) (100 ng/$\mu$L) or interferon gamma (IFN$\gamma$) (100 ng/$\mu$L) for 48 hours. B) Our 25-gene TIC signature was cross-referenced with INTERFEROME to determine which genes are regulated by interferons. C) SUM159, BCM-4272, and BCM-15006 cells were treated with interferon alpha (IFN$\alpha$) (100 ng/$\mu$L) or interferon gamma (IFN$\gamma$) (100 ng/$\mu$L) for 24 hours, then the expression of three upstream regulators (PARP9, PARP14, STAT1) and three downstream targets (ADAR, OAS2, and BST2) of interferon signaling were examined by qRT-PCR ($n$ = 3). ***$p < 0.001$, ****$p < 0.0001$; values are mean $\pm$ SD.

**Supplementary Table 7: Conserved genes between clusters with activation of interferon response genes.** Summary of every intersection between TIC clusters from each xenograft model.

| | SUM159/BCM-4272/BCM-15006 | BCM-4272/BCM-15006 | SUM159/BCM-4272 | SUM159/BCM-15006 |
|---|---|---|---|---|
| **1** | BST2 | OAS1 | IFI30 | NR4A2 |
| **2** | ISG20 | IFI44L | BST2 | BST2 |
| **3** | PLSCR1 | OAS2 | IRF6 | ITM2B |
| **4** | IFI6 | IFIT1 | ISG20 | ISG20 |
| **5** | OAS2 | RSAD2 | PLAAT4 | KRT15 |
| **6** | OAS3 | HERC5 | BIRC3 | SCPEP1 |
| **7** | PARP14 | BST2 | CDH3 | PLSCR1 |
| **8** | ISG15 | XAF1 | PLSCR1 | RCAN1 |
| **9** | PML | IFIT2 | BCL3 | IRF1 |
| **10** | PSME1 | IFI44 | IFI6 | IFI6 |
| **11** | OAS1 | OAS3 | OAS2 | OAS2 |
| **12** | ADAR | IFITM1 | CD47 | MCL1 |
| **13** | IFITM3 | CMPK2 | OAS3 | OAS3 |
| **14** | SHISA5 | SAMD9 | PARP14 | PARP14 |
| **15** | IFITM1 | MX1 | ARNTL2 | ISG15 |
| **16** | MT2A | ISG15 | ISG15 | NUB1 |
| **17** | PSME2 | PARP14 | GBP2 | IFI27 |
| **18** | PSMB10 | IFI6 | PSMB9 | PML |
| **19** | GNB4 | DDX60L | PML | CAPG |
| **20** | PARP9 | DDX60 | ZBTB38 | PSME1 |
| **21** | EIF2AK2 | SHFL | UBE2L6 | OAS1 |
| **22** | IFI44L | HERC6 | PSME1 | CD59 |
| **23** | SP100 | DDX58 | OAS1 | ADAR |
| **24** | STAT1 | EIF2AK2 | PSMA6 | TMEM219 |
| **25** | HLA-C | IFITM3 | INHBA | IFITM3 |
| **26** | | USP18 | ADAR | SHISA5 |
| **27** | | PARP12 | PLAUR | IFITM1 |
| **28** | | IFIH1 | IFITM3 | MT2A |
| **29** | | STAT1 | SHISA5 | PSME2 |
| **30** | | SP100 | BLVRB | PSMB10 |
| **31** | | PLSCR1 | IFITM1 | GNB4 |
| **32** | | HELZ2 | MT2A | PARP9 |
| **33** | | ZNFX1 | PSME2 | ITGA6 |
| **34** | | SP110 | PSMB10 | EIF2AK2 |
| **35** | | APOL6 | GNB4 | IFI44L |
| **36** | | GBP1 | PARP9 | SP100 |
| **37** | | PARP9 | RIPK2 | STAT1 |
| **38** | | PARP10 | TRIM22 | HLA-C |
| **39** | | GSDMD | EIF2AK2 | |
| **40** | | RNF213 | IFI44L | |
| **41** | | TRIM25 | NAP1L1 | |
| **42** | | ADAR | IL32 | |
| **43** | | SLFN5 | CFLAR | |
| **44** | | NMI | SP100 | |
| **45** | | RBCK1 | STAT1 | |
| **46** | | DTX3L | HLA-A | |
| **47** | | PNPT1 | HLA-C | |
| **48** | | IRF9 | MT-ATP8 | |
| **49** | | LY6E | | |
| **50** | | MT2A | | |
| **51** | | PML | | |
| **52** | | GNB4 | | |
| **53** | | CFB | | |
| **54** | | B2M | | |
| **55** | | ZC3HAV1 | | |
| **56** | | OPTN | | |
| **57** | | TAPBP | | |
| **58** | | TRIM14 | | |
| **59** | | PSMB10 | | |
| **60** | | SHISA5 | | |
| **61** | | STAT2 | | |

| | | |
|---|---|---|
| **62** | | PHF11 |
| **63** | | ISG20 |
| **64** | | SPATS2L |
| **65** | | TRIB1 |
| **66** | | PSME1 |
| **67** | | STX17 |
| **68** | | PSME2 |
| **69** | | FMR1 |
| **70** | | TRIM38 |
| **71** | | TRIM56 |
| **72** | | NCOA7 |
| **73** | | LAP3 |
| **74** | | TYMP |
| **75** | | NBN |
| **76** | | TMEM50A |
| **77** | | HLA-C |
| **78** | | HLA-B |
| **79** | | HLA-E |

**Supplementary Table 8: Our TIC gene signature derived from scRNA-seq is fundamentally different from TIC signatures derived from bulk RNA sequencing.** A comparison of the overlaps of each TIC cluster and six previously published TIC signatures. An orange highlighted box indicates a statistically significant overlap, as determined by Fisher's exact test.

| | **SUM159 C7/4272 C7: 48 genes** | **SUM159 C7/15006 C6: 38 genes** | **4272 C7/15006 C6: 79 genes** | **Creighton et al. (2009): 154 genes** | **Pei et al. (2020): 32 genes** | **Pece et al. (2019): 20 genes** | **Charafe-Jauffret et al. (2009): 28 genes** | **Leth-Larsen et al. (2012): 31 genes** | **Colacino et al. (2018) 96 genes** |
|---|---|---|---|---|---|---|---|---|---|
| **SUM159 C7/4272 C7: 48 genes** | | 25 genes p=2.9e-62 | 25 genes p=1.3e-51 | 0 genes | 0 genes | 0 genes | 0 genes | 0 genes | 1 gene |
| **SUM159 C7/15006 C6: 38 genes** | | | 25 genes p=2.3e-55 | 0 genes | 0 genes | 0 genes | 0 genes | 0 genes | 2 genes |
| **4272 C7/15006 C6: 79 genes** | | | | 0 genes | 0 genes | 1 gene | 0 genes | 0 genes | 0 genes |
| **Creighton et al. (2009): 154 genes** | | | | | 0 genes | 0 genes | 1 gene | 0 genes | 1 gene |
| **Pei et al. (2020): 32 genes** | | | | | | 0 genes | 0 genes | 0 genes | 0 genes |
| **Pece et al. (2019): 20 genes** | | | | | | | 0 genes | 0 genes | 0 genes |
| **Charafe-Jauffret et al. (2009): 28 genes** | | | | | | | | 0 genes | 1 gene |
| **Leth-Larsen et al. (2012): 31 genes** | | | | | | | | | 1 gene |
| **Colacino et al. (2018) 96 genes** | | | | | | | | | |

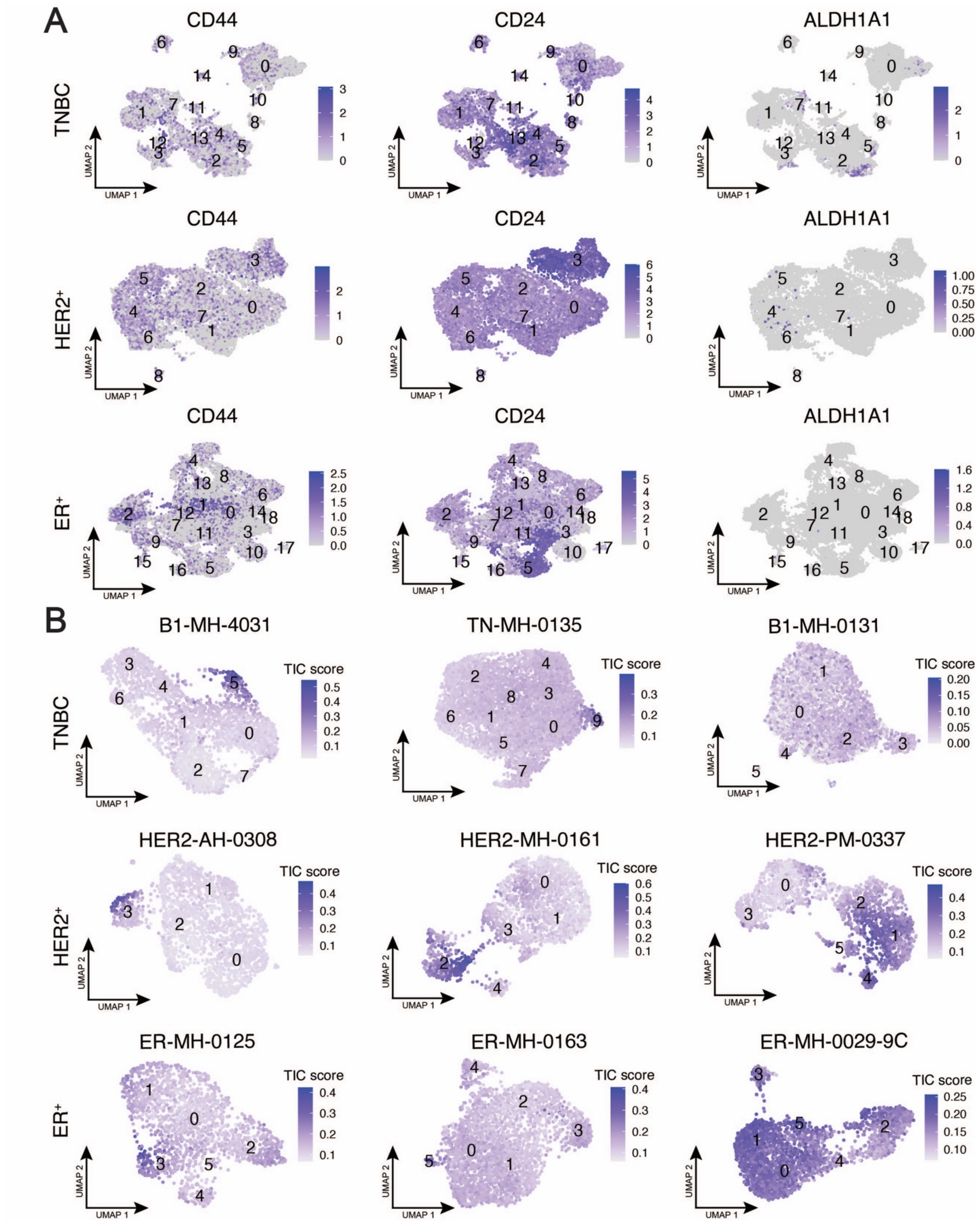

**Supplementary Figure 7: Interferon response genes are differentially expressed in a subset of epithelial cells in human breast cancer patient samples.** A) Feature plots of the expression of TIC markers CD44, CD24, and ALDH1A1 in TNBC, $HER2^+$, and $ER^+$ breast cancer patients.

B) TIC score feature plots of three individual patients of each breast cancer subtype.

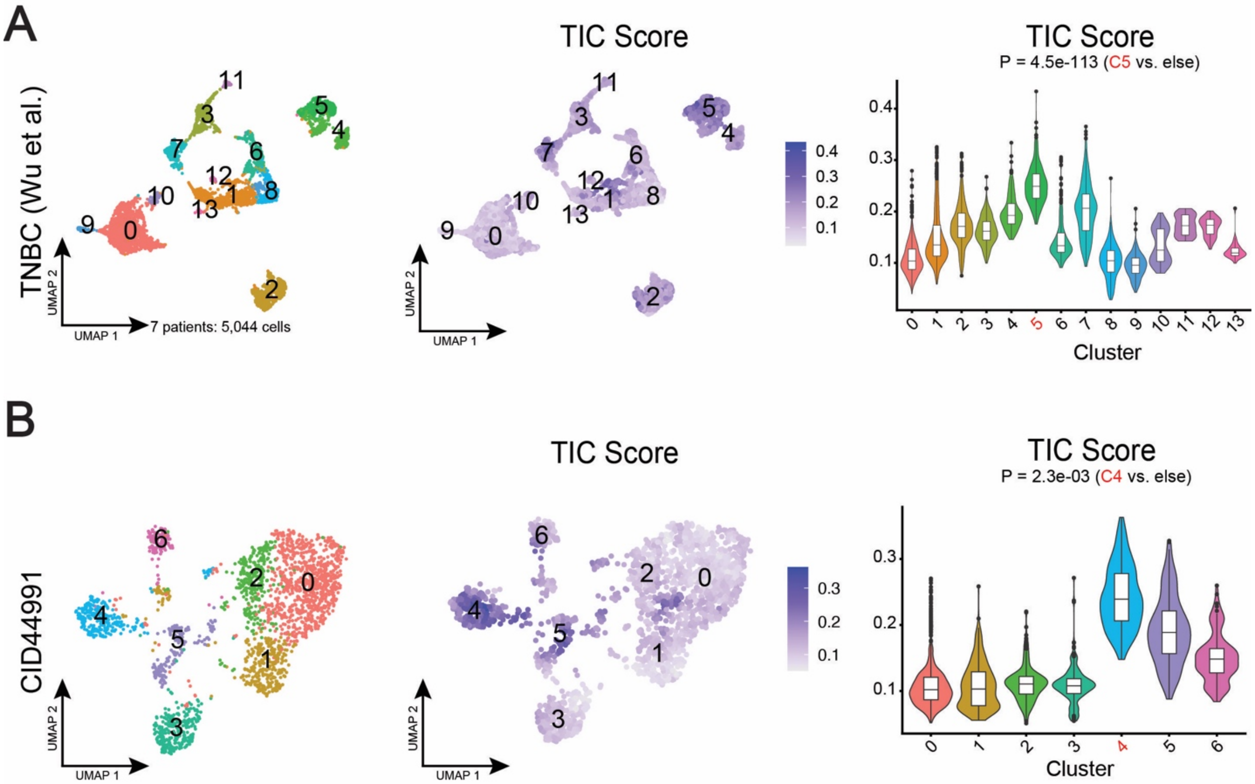

**Supplementary Figure 8: Interferon response genes are differentially expressed in a subset of epithelial cells in human breast cancer patient samples.** A) UMAP visualization of seven human TNBC patient samples with feature plot and violin plot showing TIC score, derived from based on the expression of our 25-gene TIC signature. A two-sided Wilcoxon test was used to determine significance of TIC score. B) UMAP visualization of the only human TNBC patient sample in the Wu et al. dataset with >1,000 non-cycling epithelial cells, a TIC score feature plot, and violin plot showing TIC score, derived from based on the expression of our 25-gene TIC signature. A two-sided Wilcoxon test was used to determine significance of TIC score.
